# Archaic translation initiation factor eIF5B supports KSHV late lytic replication and viral oncogenesis by mimicking a hypoxic cellular landscape

**DOI:** 10.64898/2026.02.23.707408

**Authors:** Christian McDonald, Omayra Méndez-Solís, Yuan-De Tan, Renu Khasa, Julian Naipauer, Anuj Ahuja, Efe Karaca, Yoandy Marcelo, Natalie Ruiz-Ocaña, Clarice Djouwoug Noussi, Thai My Duyen Tran, Joshua M Hare, Sabita Roy, Stephen Lee, Enrique A. Mesri, Noula Shembade

## Abstract

Kaposi’s Sarcoma (KS) Herpesvirus (KSHV) is the etiological agent of KS, an AIDS-defining illness. Recent studies have demonstrated that even during normoxic conditions, KSHV facilitates replication by modulating hypoxia-inducible factors (HIFs), creating a hypoxia-like environment that promotes cellular transformation. KSHV lytic viral genes are favored for protein synthesis during infection by upregulating HIF2α and utilizing the hypoxic eIF4E2 translation initiation complex in oxygen-replete conditions. This translation initiation plasticity (TRIP) links viral replication strategies to angiogenic signaling and oncogenicity. However, the molecular basis of this plasticity remains poorly understood. This study reveals that viral inhibition of eukaryotic initiation factor 2 (eIF2) induces the use of another alternative initiation factor, eIF5B, which canonically mediates delivery of initiator methionine-tRNAi^Met^ during hypoxia. We demonstrate through ribosome density fractionation that eIF2 accumulates in translationally inactive monosome fractions while eIF5B redistributes its translation activity toward polysomes during lytic replication. Progressive dependence on eIF5B during KSHV infection was illustrated by impaired late lytic gene expression, diminished virion production, and altered polysome profiles following eIF5B knockdown. Transcriptomic analyses further reveal that the mRNA landscape in KSHV-infected cells depleted of eIF5B mirrors that of uninfected hypoxic cells. Moreover, silencing of eIF5B in a natural infection model reduces both VEGF secretion and anchorage-independent growth—two hallmarks of KSHV viral oncogenicity. These results demonstrate that eIF5B functions as an essential component of alternative translation initiation machinery activated in response to eIF2 inactivation during KSHV lytic replication, emphasizing its key role in viral pathogenesis and its potential as a novel therapeutic target.

**IMPORTANCE:** Kaposi’s Sarcoma Herpesvirus (KSHV) is a human cancer virus that causes severe malignancies in immunocompromised individuals, including the blood vessel cancer Kaposi’s Sarcoma and a highly aggressive body-cavity lymphoma. Although cells mount a response to infection by shutting down global protein synthesis, how viruses like KSHV continue making viral proteins when this cellular machinery is shutdown remains unclear. This study uncovers that KSHV exploits an ancient cellular factor typically used for protein production under low-oxygen conditions. We identify a previously unrecognized role for the translation factor, eIF5B, in supporting KSHV replication and select production of viral proteins during infection. Our work further demonstrates that KSHV’s utilization of eIF5B contributes to a hypoxia-like environment during infection, ultimately contributing to cancer-promoting changes within the cell. These findings highlight a new strategy for tumor-virus reprogramming of host cells, opening an avenue for novel therapeutic targets to eliminate virus-associated cancers.

## INTRODUCTION

Viral-mediated cellular transformation can activate several hallmarks of cancer and is responsible for approximately 12-15% of human malignancies worldwide^1,2^. Kaposi’s Sarcoma (KS) herpesvirus (KSHV) is one of the seven known oncogenic viruses in humans^1^. Along with epithelial cells, this γ-herpesvirus has cellular tropisms for endothelial cells, B cells, mesenchymal cells and fibroblast cells^1,3,4–6^. Because of its broad cellular host range, KSHV has been identified as the etiological agent of several malignancies, most notably the aggressive B-cell Primary effusion lymphoma and KS, both AIDS-defining cancers^7,8^. Although AIDS-KS has seen a decline since the widespread use of antiretroviral therapies, persistent KS remains a significant clinical challenge, with complete disease remission achieved in approximately only 50% of patients^9^.

Viruses are obligate intracellular pathogens that hijack host cellular machinery for replication, demanding a fine balance between exploiting host resources and deploying strategies to subvert cellular defenses that could cripple virion production^10^. Specifically, a salient feature of viral propagation is the dependence on host protein synthesis machinery^10,11^. To initiate eukaryotic translation, 12 core eukaryotic initiation factors (eIFs) composed of over 30 polypeptide subunits are required^12,13^. A subset of these translation factors plays a key role in the formation of the eIF4F initiation complex. These proteins include the cap-binding protein eIF4E, the eIF4G scaffold protein and the eIF4A RNA helicase, which have all been shown to be co-opted by viruses during infection^11,14^. In contrast to RNA viruses, double-stranded DNA viruses, like KSHV, predominantly commandeer the canonical cap-mediated mode of translation^11,15^, although cap-independent mechanisms have been observed^16,17^.

Since protein synthesis is a metabolically intensive process, the initiation step serves as a crucial point in translational regulation, especially during cellular stresses such as hypoxia. Cells can adapt to hypoxic environments through the stabilization of heterodimeric hypoxia-inducible factors (HIFs)^18,19^. Discovery of the HIFs led to a prevailing model where HIF-directed transcriptional changes were at the center of the cellular response to oxygen deficiency^18–20^. However, understanding of an adaptive translational paradigm during hypoxia has shifted this framework^21^. Earlier studies suggested that inhibition of the canonical eIF4F complex under hypoxic conditions resulted in a widespread decrease in cap-dependent translation and, therefore, a global decline in protein synthesis ^22–25^. Instead, we now understand that under hypoxic stress, cells recruit an alternate and mTOR-independent machinery known as the eIF4F^H^ complex^26,27^. Composed of HIF2α, the cap-binding protein eIF4E2, scaffolding protein eIF4G3 and RBM4^26,27^, amongst other RNA-binding proteins^28^. This alternate complex allows for a select pool of mRNAs to undergo protein synthesis during hypoxic stress^28–29^.

Interestingly, patients with KS present with lesions throughout the body, but are prevalent in hypoxic regions, including the skin of the lower extremities^30,31^. The potential correlation between hypoxia and the development of KS has prompted research indicating that hypoxia promotes the replication of KSHV, while KSHV simultaneously upregulates HIF1α, producing a positive feedback loop that induces a hypoxia-like environment in infected cells^30,32,19,32,33^. Recent work has demonstrated that during lytic KSHV infection in normoxia, upregulation of HIF2α leads to the synthesis of lytic viral mRNAs through the cap-binding protein eIF4E2 for translation initiation^34–35^, providing an intriguing viral strategy to optimize viral gene expression in the host cell. This interplay underscores the importance of the hypoxic cellular machinery during KSHV replication and oncogenesis.

Reduction in global translation is a major response to virtually all classes of cellular stresses in order to suppress the synthesis of nonessential proteins and prevent further energy depletion in the cell^36^. One of the most established mechanisms for achieving this includes the phosphorylation of eIF2⍺ at the level of translation initiation. The eIF2 heterotrimeric protein plays a crucial role in anchoring the initiator methionine-tRNAi^Met^ to the 40S ribosomal subunit during initiation of translation^10,37^. Phosphorylation of the alpha (⍺) subunit at Ser51 is mediated by one of the four mammalian protein kinases that respond to discrete types of stresses^11,36,38^. During environmental stresses like hypoxia, it was previously thought that this kinase-mediated phosphorylation would inhibit eIF2 activity and abolish translation initiation entirely. However, the observations of robust protein synthesis under hypoxic conditions via the eIF4F^H^-mediated translation machinery led to the discovery that the initiation factor eIF5B preferentially functions in hypoxia to deliver the initiator methionine-tRNAi^Met^ following eIF2 inactivation^39,40^.

To sustain primordial life in anoxic environments, the ancient initiation factor-2 (IF2), was used to catalyze crucial reactions essential for early life in these oxygen-deficient conditions^41^. This GTPase may be traced back to the last universal common ancestor (LUCA) and is evolutionarily conserved in eukaryotic organisms as eIF5B^42,43^. In modern eukaryotes, eIF5B is largely dispensable under normoxic conditions and primarily facilitates the late-stage joining of ribosomal subunits during translation initiation, highlighting its homology with the prokaryotic IF2 ^37,38,42,44^. Considering the evidence that eIF5B activity can functionally substitute for eIF2 in hypoxic cells and recognizing KSHV’s dependence on hypoxic-adapted cellular mechanisms, we hypothesized that KSHV may preferentially recruit eIF5B for replication and evasion of host stress responses.

In this study, we report that during KSHV lytic infection, eIF2 inhibition leads to increased expression of eIF5B, which is specifically important in the synthesis of late lytic mRNAs and virion production. Depletion of eIF5B also led to global changes in the mRNA landscape, mimicking hypoxic cell environment and ultimately contributing to KSHV-viral oncogenic mechanisms.

## RESULTS

### KSHV lytic reactivation leads to eIF2 inhibition and increased eIF5B expression

During lytic KSHV infection, upregulation of HIF2α leads to the translation of lytic viral mRNAs through eIF4E2^34^. In hypoxia, this translation-initiation complex has a weak affinity for eIF2 but preferentially utilizes the hypoxic surrogate, eIF5B (**Figure 1A**)^39,40^. Thus, we hypothesized that since KSHV exploits the hypoxic translation initiation machinery under normoxic conditions, it may favor the recruitment of eIF5B for replication and viral protein synthesis.

**Figure 1:**
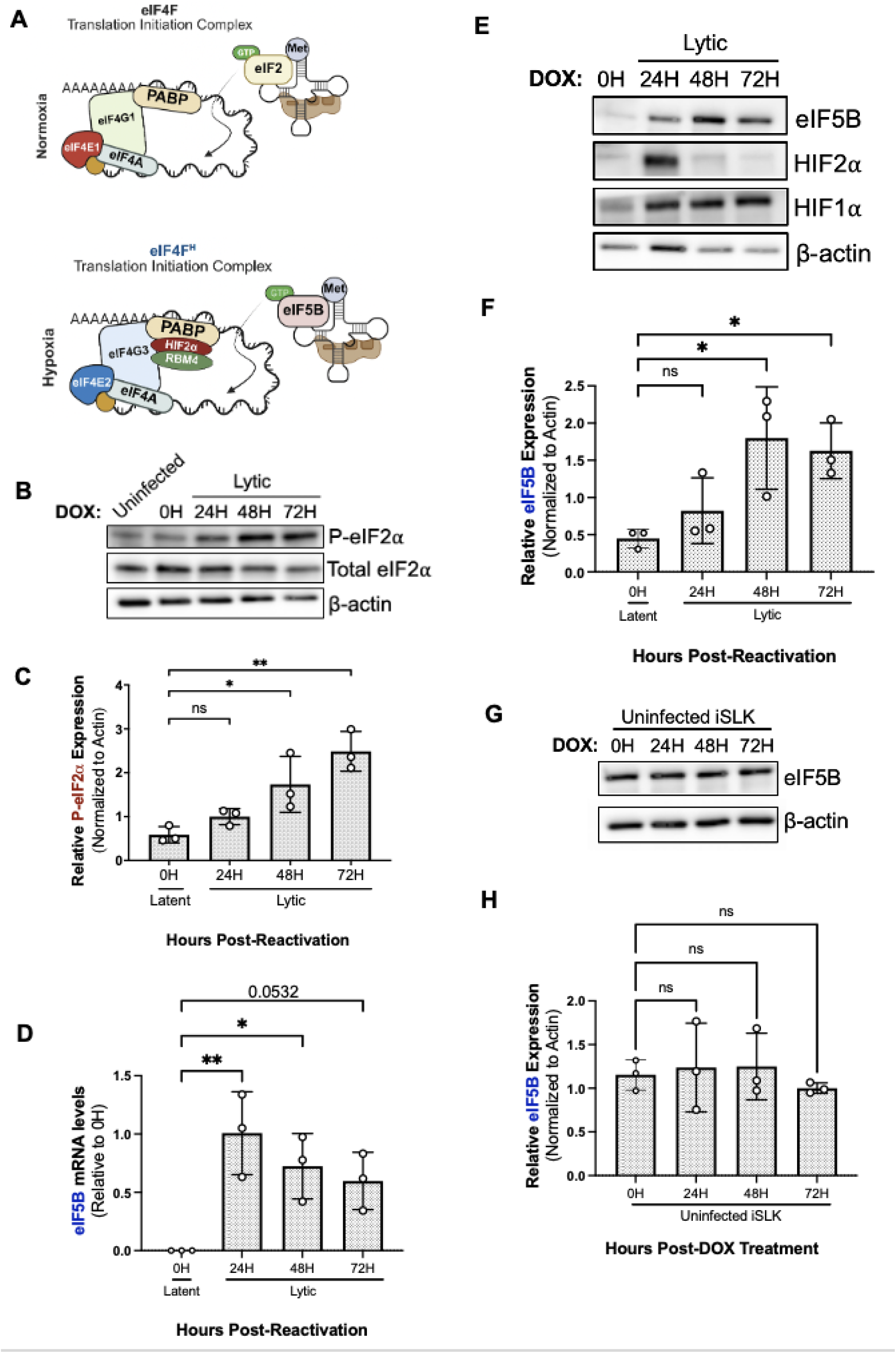
KSHV lytic reactivation leads to eIF2 inhibition and increased eIF5B expression. **(A)** Schematic diagram of the canonical eIF4F translation inititation complex and the alterantive eIF4F^H^ complex used by cells under hypoxic stress. **(B)** Representative immunoblot of eIF2 and P-eIF2a protein levels in uninfected iSLKs, before (0H) and after (24-72H) DOX treatment in iSLK.219 cells cultured in normoxia. **(C)** Quantification of (A) calculated using Image J (n=3; mean ± SD; **p < 0.0231, ***p < 0.0013, One-Way ANOVA with Dunnett’s Multiple Comparison Test). **(D)** eIF5B mRNA levels in iSLK.219 cells before (0H) and after (24-72H) DOX treatment in normoxia relative to 0H, measured by qRT-PCR (n=3, mean ± SD; *p < 0.00221, **p < 0.0036, One-Way ANOVA with Dunnett’s Multiple Comparison Test. **(E)** Representative immunoblot of eIF5B, HIF1a and HIF2a protein levels before (0H) and after (24-72H) DOX treatment in iSLK.219 cells cultured in normoxia. **(F)** Quantification of (E) calculated using Image J (n=3; mean ± SD; *p < 0.0165, **p < 0.0321, One-Way ANOVA with Dunnett’s Multiple Comparison Test). **(G)** Representative immunoblot of eIF5B protein levels in uninfected iSLKs, before (0H) and after (24-72H) DOX treatment cultured in normoxia. **(H)** Quantification of (F) calculated using Image J (n=3; mean ± SD; ns= no significance, One-Way ANOVA with Dunnett’s Multiple Comparison Test)

To investigate how eIF2 and eIF5B are impacted during KSHV infection, we used the doxycycline (DOX)-inducible KSHV producer cell line, iSLK.KSHV219 (iSLK.219). This iSLK cell line harbors a recombinant virus (rKSHV219) that produces infectious KSHV virions at 48-72 hours post-reactivation^45^. To determine the impact of KSHV on these translation factors, we evaluated their expression over the course of 72 hours after viral reactivation as described previously^34^. Following lytic induction, we observed phosphorylation of the eIF2α subunit significantly increased as compared to uninfected and latently infected cells (**Figure 1B & 1C**). This increase in eIF2α subunit phosphorylation is likely due to KSHV-induced inhibition of eIF2, which coincides with the initiation of viral reactivation and replication^45^. Following this, we examined whether the expression of eIF5B is affected during doxycycline-induced KSHV replication. Interestingly, both eIF5B mRNA (**Figure 1D**) and protein levels were significantly elevated (**Figures 1E & 1F**) during lytic reactivation in comparison to uninfected iSLK cells (**Figures 1G & 1H**) and hypoxic cells (**Figure S1A**). The expression of eIF5B was significantly increased at 48- and 72-hours post-reactivation (**Figures 1E & 1F**), coinciding with the production of infectious KSHV virions suggesting a significant change of eIF5B in the late lytic stage of infection. Consistent with prior findings, we also observed an increase in HIF1α and HIF2α **(Figure 1E)**^34^.

Recent studies have demonstrated that herpesviruses, including human cytomegalovirus (HCMV), alter host eukaryotic initiation factor (eIF) accumulation^46^. We next sought to determine if the increase in eIF5B expression during KSHV infection was observed across eIFs or specific to this translation factor. Evaluation of eIFs specific to the canonical eIF4F complex (eIF4E1, eIF4G1 and eIF4A) as well as the hypoxic translation initiation eIF4F^H^ complex (eIF4G3 and RMB4) revealed no augmented changes in protein levels comparable to eIF5B during KSHV lytic reactivation (**Figures S1B-D**). Interestingly, the expression levels of helicase eIF4A, RNA-binding protein RBM4 and scaffolding protein eIF4G3 were all decreased during KSHV lytic replication (**Figures S1C & S1D**).

The phosphorylation of eIF2 during KSHV infection (**Figures 1A & 1B**) suggests the need for an eIF2-independent translation factor to assist viral replication during this period of eIF2 inactivation. Inhibition of eIF2 over 72 hours of KSHV reactivation parallels the eIF5B-specific increase in transcript abundance and gene expression, pointing to a possible molecular shift towards using eIF5B that would allow for viral replication.

### eIF5B plays a critical role in KSHV lytic replication and selective viral gene expression

Prior research has indicated that under normoxic conditions, KSHV recruits the eIF4F^H^ complex, with two of its components, HIF2α and eIF4E2, being essential for optimal viral replication^34^. Given the increase in eIF5B mRNA and protein levels (**Figures 1D-F**), we next sought to determine if eIF5B is required for productive KSHV replication during normoxia by using the DOX-inducible iSLK.219 cellular model. This cell line expresses GFP (latent infection marker) under the control of the human constitutive EF1-α promoter, and RFP (lytic and reactivation marker) regulated by the KSHV lytic PAN promoter, which leads to the generation of new infectious viral particles 48-72 hours after DOX treatment^45^. To determine if eIF5B plays a role in KSHV lytic replication, iSLK.219 cells transfected with non-silencing, control siRNA or eIF5B siRNA for 24 hours were treated with doxycycline to induce KSHV lytic reactivation **(Figure 2A)**. We observed approximately 50% reduction in KSHV lytic reactivation in cells treated with eIF5B siRNA compared to control siRNA-treated cells (**Figures 2B & 2C**) as determined by RFP expression 48 hours after lytic induction (**Figures S2A**)

**Figure 2:**
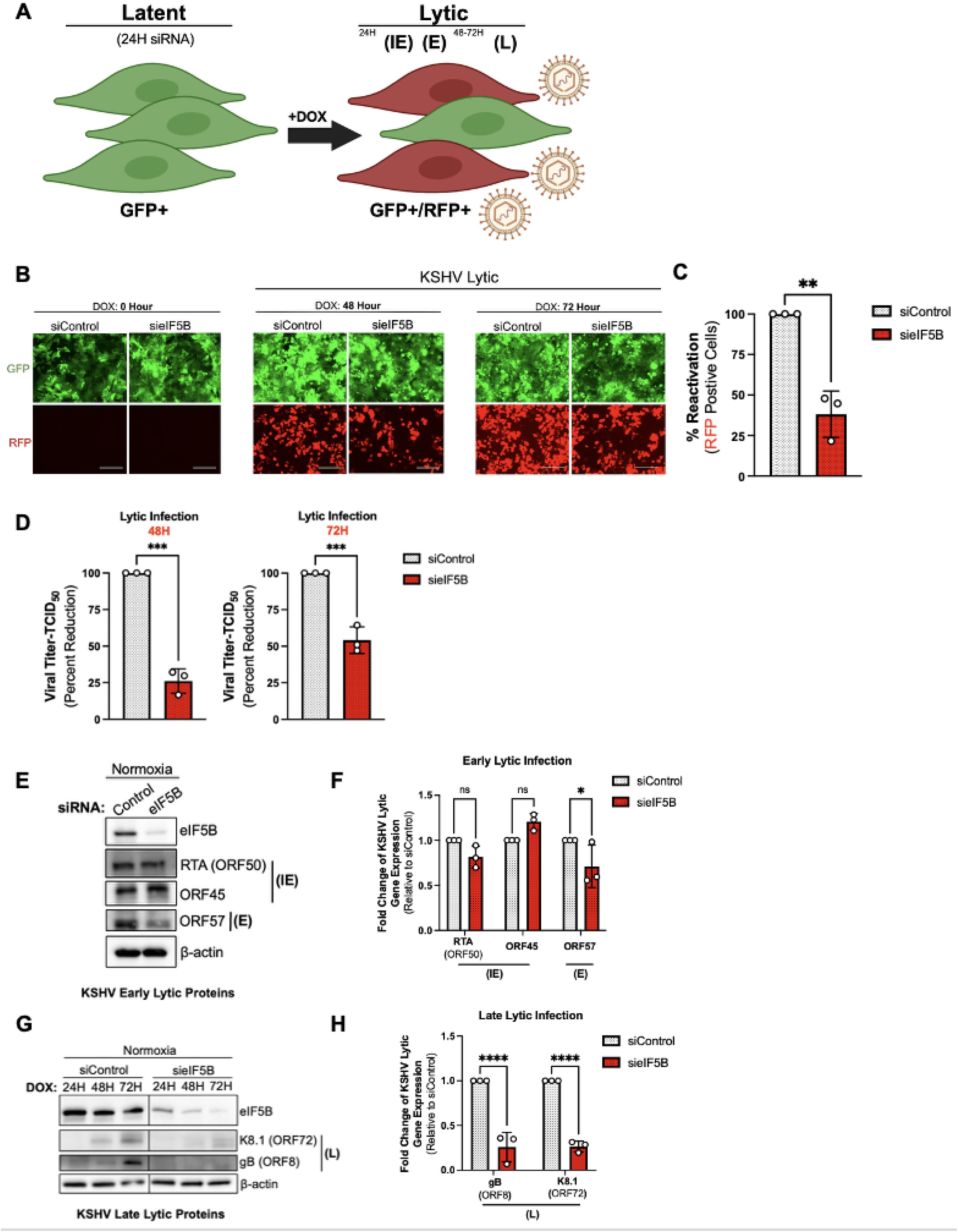
eIF5B plays critical role in KSHV lytic replication and selective viral gene expression. **(A)** Schematic diagram of iSLK.219 infection system. These cells express GFP during latent infection and upon the addition of doxycycline (DOX), these cells express the reactivation marker RFP, driven by the KSHV lytic PAN promoter with progress increase in the immediate-early (IE), early (E) and late lytic (L) viral protein levels as well as virion production. **(B)** Images of iSLK.219 cells were silenced for 24 hours. Images indicate before (0H) and after (24-72H) DOX treatment in normoxia. Scale bar 100mm. **(C)** Percentage of reactivation 48H post-dox of sieIF5B relative to siControl. RFP expression was considered the reactivation marker and was measured using flow cytometry. **(D)** Infectious virion production of siControl and sieIF5B cells post-reactivation measured by HEK-AD293 TCID_50_ (n=3; mean ± SD; 48H: ***p < 0.0001, 72H:***p < 0.0009, Unpaired t-test) **(E)** Representative immunoblot of KSHV immediate-early (IE) and early (E) protein levels in siControl and sieIF5B silenced cells 48H after DOX induction. **(F)** Quantification of (E) calculated using Image J. **(G)** Representative immunoblot of KSHV late lytic (L) protein levels in siControl and sieIF5B silenced cells after (24-72H) DOX treatment cultured in normoxia. **(H)** Quantification of (G) calculated using Image J

Next, we investigated the effect of eIF5B silencing on infectious virus production by harvesting cell-free supernatants from non-silenced and eIF5B-depleted cells and used them to determine the 50% tissue culture infective dose (TCID_50_). Transient silencing of eIF5B markedly reduced the production of infectious virions at both 48 and 72 hours after lytic induction in normoxia by 74% or 46%, respectively (**Figure 2D & S2B**). These observations were not attributed to sieIF5B-associated suppression of cell growth and viability (**Figures S2C-E)** or reduction in DOX-inducible RTA mRNA abundance **(Figure S2F).** This is consistent with previously published findings that eIF5B is non-essential in normoxia^39,43,47^.

The iSLK.219 cells provide a platform to study the KSHV latent-lytic switch which results in the temporally regulated cascade of viral gene expression: immediate early (IE), early (E) and late lytic (L) KSHV genes (**Figure 2A**). Leveraging this system, we next evaluated the impact of eIF5B depletion on lytic viral protein synthesis by immunoblot. Interestingly, lytic viral translation of the immediate-early proteins RTA (ORF50) and ORF45 remained largely unchanged (**Figures 2E & 2F**). In contrast, as the KSHV lytic cycle progressed, we observed a reduction of early lytic ORF57 along with a near-complete ablation of late lytic viral proteins, gB (ORF8) and K8.1 (**Figures 2G-H**). Additionally, we investigated eIF5B’s role during KSHV latency (**Figure S3**). eIF5B silencing did not affect KSHV latent infection or latent gene expression (**Figure S3**). These observations strengthen the importance of eIF5B as a crucial translation factor during the expression of late lytic viral genes.

### eIF5B concentrates in translating ribosomes during KSHV infection

To investigate the translational role of eIF5B during KSHV lytic replication, we employed the use of ribosome density fractionation, which measures translational activity by separating mRNAs in a sucrose gradient based on the density of ribosomes bound to transcripts (**Figure 3A**). Transcripts that are disengaged from protein synthesis, with less ribosomes, migrate to the top portion of the sucrose gradient following centrifugation (**Figure 3A**). Actively translating transcripts are bound by several ribosomes, known as polysomes, which are consigned to the bottom of the gradient (**Figure 3A**).

**Figure 3:**
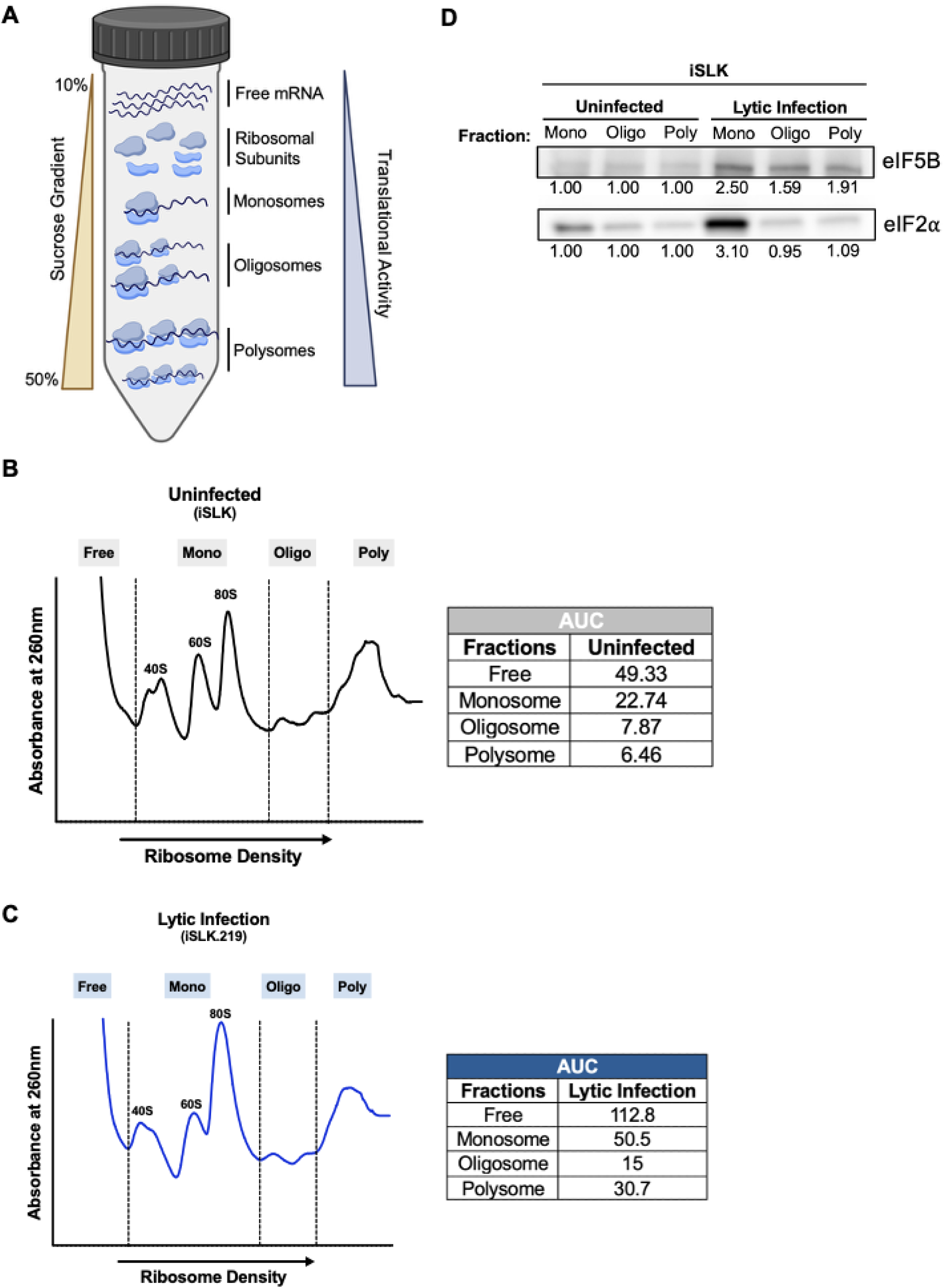
eIF5B concentrates in translating ribosomes during KSHV infection. **(A)** Schematic of the ribosoome density fractionation technique which enables seperation of mRNAs based on number of bound ribosomes and subunuts**. (B)** Ribosome density profiles of uninfected iSLK cells following addition of DOX for 48 hours. Cells were treated with cyclohexamide and lysates were sedimented through sucrose gradients and fractionated. Ribosomal subunits (40S and 60S), monosomes (80S), oligosomes (oligo) and polysomes (poly) are indicated. The table shows the area under the curve (AUC) for each fraction. The table shows the area under the curve (AUC) for each fraction. **(C)** Ribosome density profiles of reactivated iSLK.219 cells following addition of DOX for 48 hours. The table shows the area under the curve (AUC) for each fraction. **(D)** Immunoblot of uninfected iSLK and reactivated iSLK.219 after the addition of DOX for 48 hours. Fractions: Monosome (Mono), Oligosome (Oligo) and Polysome (Poly). Equal volumes of lysate was loaded in all lanes.

To perform ribosome density fractionation, we first added DOX to both uninfected iSLK cells and KSHV-infected iSLK.219 cells for 48 hours, followed by cycloheximide treatment to terminate translation elongation. Ribosome density profiles of lytically induced iSLK.219 cells exhibited a significant shift compared to KSHV-negative iSLK cells (**Figures 3B & 3C**). Notably, the polysome profile of KSHV-infected cells resulted in a substantial increase in translating ribosomes, as determined by quantitative analysis of the area under the curve (**Figure 3C**).

To evaluate the translation efficiency of canonical eIF2 and eIF5B after lytic reactivation, protein lysates from each fraction were subjected to immunoblot analysis using anti-eIF5B and anti- eIF2 antibodies. Remarkably, examination of these fractions revealed increased polysome association of eIF5B, while eIF2 disengaged from polysomes during KSHV infection compared to uninfected iSLK cells (**Figure 3D**). This trend of eIF2 protein levels to monosome fractions in contrast to eIF5B-enriched polysome fractions suggests an important change in the translational role of eIF5B during KSHV infection in normoxia.

### Depletion of eIF5B reduces global protein synthesis and depletes viral mRNAs from polysomes

Findings from Figures 1-3 indicate that eIF5B plays a key role in viral protein synthesis during lytic replication. We next examined the effect of eIF5B knockdown on translation initiation during KSHV infection. First, no significant changes were observed in ribosome density profiles between control and eIF5B-silenced groups in uninfected iSLK cells after DOX induction (**Figure 4A & S5A**). Strikingly, knockdown of eIF5B in lytically induced iSLK.219 cells had a substantial effect on global mRNA translation activity (**Figure 4B**), highlighted by the shifts in ribosome density profiles in both the oligosome (mild translation) and polysome (active translation) fractions (**Figure 4B**).

**Figure 4:**
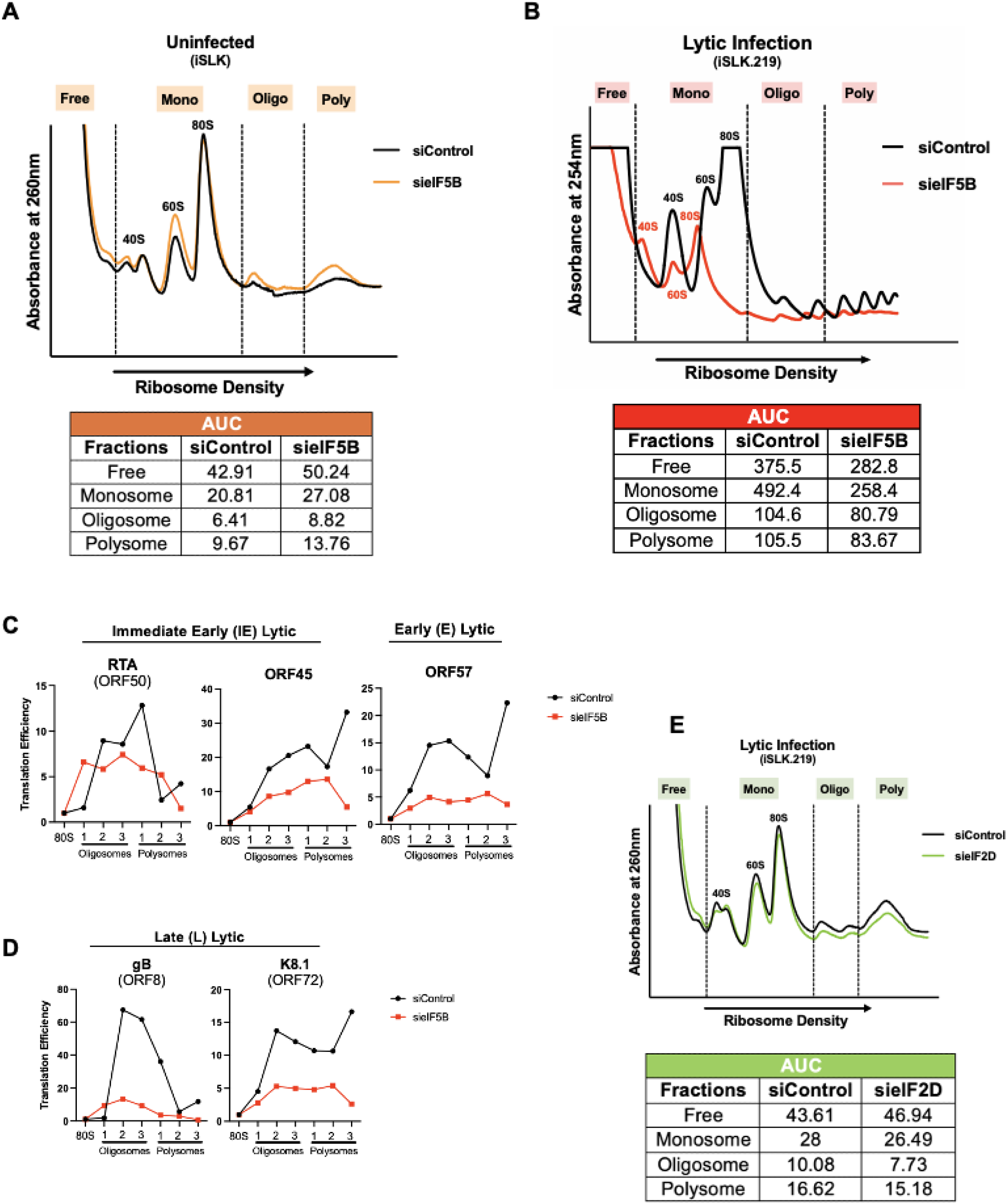
Depletion of eIF5B reduces global protein synthesis and depletes viral mRNAs from polysomes. **(A)** Ribosome density profiles of uninfected iSLK cells silenced (siControl or sieIF5B) for 24 hours followed by the addition of DOX for 48 hours. Silenced cells were treated with cyclohexamide and lysates were sedimented through sucrose gradients and fractionated. Ribosomal subunits (40S and 60S), monosomes (80S), oligosomes (oligo) and polysomes (poly) are indicated.The table shows the area under the curve (AUC) for each fraction. **(B)** Ribosome density profiles of iSLK.219 cells silenced (siControl or sieIF5B) for 24 hours followed by the addition of DOX for 48 hours to induce KSHV lytic replication. Silenced cells were treated with cyclohexamide and lysates were sedimented through sucrose gradients and fractionated. Ribosomal subunits (40S and 60S), monosomes (80S), oligosomes (oligo) and polysomes (poly) are indicated. The table shows the area under the curve (AUC) for each fraction. **(C)** Translation efficiency of KSHV (IE) and (E) lytic mRNAs from (B). KSHV viral mRNAs were detected using qRT-PCR (n=3). Each fraction CT value was normalized to the 80S fraction CT value. **(D)** Translation efficiency of KSHV (L) lytic mRNAs from (B). KSHV viral mRNAs were detected using qRT-PCR (n=3). Each fraction CT value was normalized to the 80S fraction CT value. **(E)** Ribosome density profiles of iSLK.219 cells silenced (siControl or sieIF2D) for 24 hours followed by the addition of DOX for 48 hours to induce KSHV lytic replication. Silenced cells were treated with cyclohexamide and lysates were sedimented through sucrose gradients and fractionated. Ribosomal subunits (40S and 60S), monosomes (80S), oligosomes (oligo) and polysomes (poly) are indicated.The table shows the area under the curve (AUC) for each fraction.

To further dissect the translational role of eIF5B on KSHV genes, RNA from each fraction was isolated and viral genes were measured by qRT-PCR. Depletion of eIF5B resulted in diminished translation efficiency of KSHV early (E) and late (L) genes, whereas the impact on immediate-early (IE) genes such as KSHV RTA was less pronounced (**Figures 4C & 4D**). The observed displacement of KSHV E and L gene transcripts from polysome fractions following eIF5B silencing is consistent with previous findings of decreased viral gene expression of these proteins in eIF5B-depleted cells (**Figure 3**). These results suggest that eIF5B is critical for viral protein synthesis of KSHV E and L genes in normoxia.

Similar to eIF5B, eukaryotic translation initiation factor eIF2D can deliver of met-tRNAi^Met^ ^48,49^. To test whether KSHV may be using other alternate eIFs during lytic replication, we silenced eIF2D and performed ribosome density fractionation in lytically reactivated iSLK.219 cells (**Figure S5B**). Knockdown of eIF2D showed no measurable effect on the ribosome density profiles in KSHV-infected cells (**Figure 4E**). Moreover, silencing eIF5B in latently infected iSLK.219 cells did not significantly alter polysome profiles (**Figure S4C**), nor was it associated with reduction of global protein synthesis by other gammaherpesviruses (**Figure S5**). These findings indicate that KSHV-infected iSLK cells exhibit increased sensitivity to eIF5B silencing relative to other cells, suggesting that eIF5B may functionally substitute for eIF2 following KSHV-induced suppression of eIF2 during lytic reactivation

### Interrogation of the global host mRNA landscape in KSHV-infected cells by eIF5B

Next, we sought to examine eIF5B-dependent host mRNA targets during lytic reactivation by performing bulk-RNA sequencing. KSHV-harboring iSLK.219 cells were transfected with either control or eIF5B siRNA for 24 hours, followed by lytic induction and collection of mRNAs (**Figure 5A**). In both groups we generated three biological replicates over the course of KSHV infection. Here, we focus on 48 hours post-reactivation because of the pronounced differences in eIF5B expression and effects on KSHV biology observed previously (**Figures 2 & 3**). Transcriptional profiles of non-silencing and eIF5B-depleted cells during KSHV reactivation were compared and identified a robust set of differentially expressed genes (1,378 genes were downregulated and 2,046 genes upregulated) **(Figure 5B).** Pathway enrichment analysis uncovered a significant overrepresentation of biological processes involved in immune response and signaling, including interferon gamma response, interferon alpha response and TNF signaling (**Figure 5C**). We confirmed this by evaluating through qPCR differentially expressed genes which are involved in these innate immune responses to viral infection, such as ISG15, IFIT3 and IL6^50,51^ (**Figure S6**). Because eIF2α is a target of host-stress and innate immunity regulation^36,52^, these findings implicate that use of eIF5B during eIF2 inhibition in normoxia may confer benefits to KSHV infection (**Figure 5B-C**). A notable observation was the mRNAs significantly enriched in pathways including hypoxia and glycolysis (**Figure 5B and S6)**. The interplay between hypoxia and KSHV infection has been well established^53^, however, no research has previously described the interaction between eIF5B and KSHV lytic replication. Because eIF5B has only been shown to replace eIF2 in hypoxia, this data supports an intriguing narrative where KSHV can upregulate eIF5B for replication and protein synthesis in normoxia.

**Figure 5:**
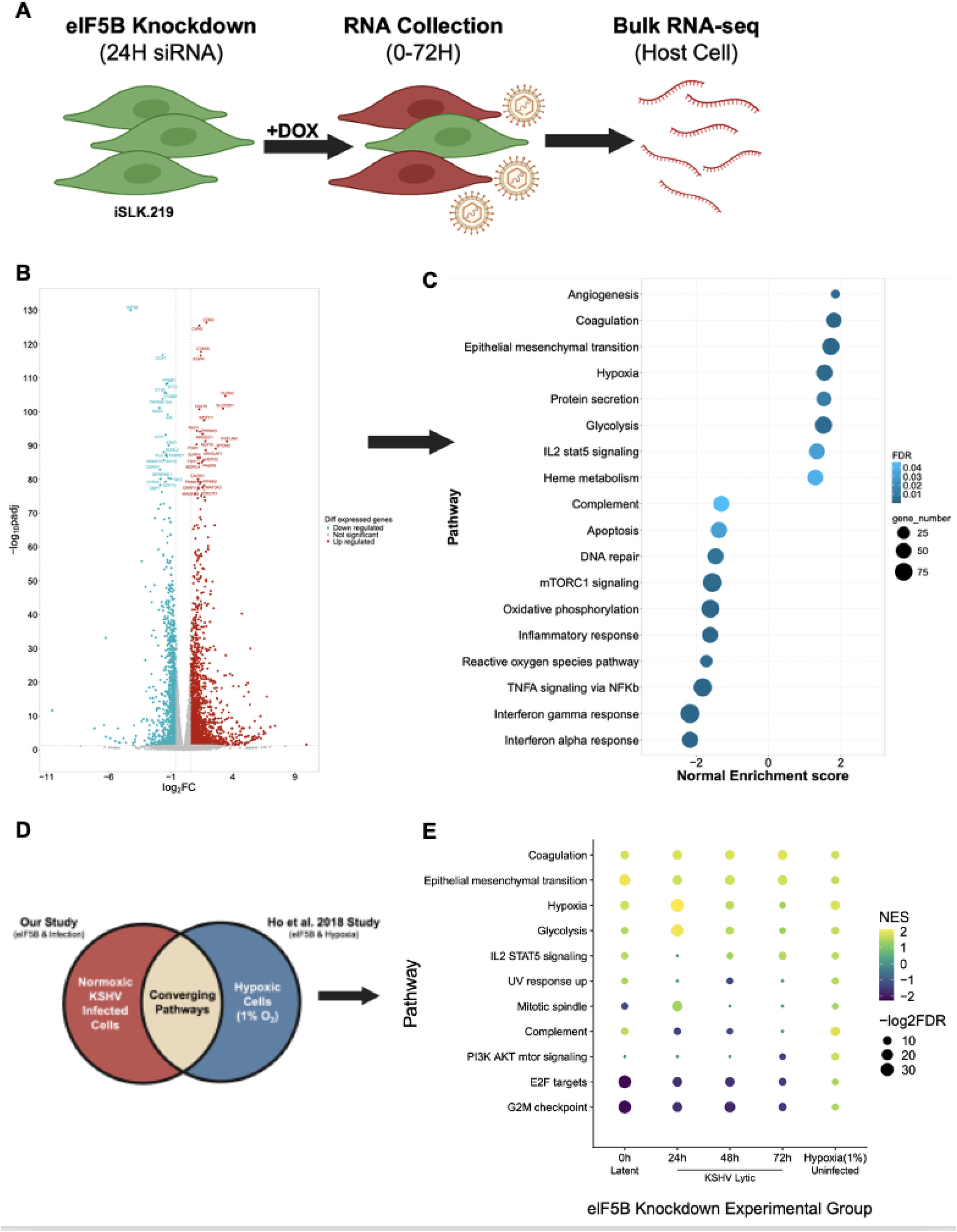
Interrogation of global host mRNA landscape in KSHV-infected cells by eIF5B. **(A)** Schematic of experimental design for Bulk-RNAseq study in eIF5B depleted KSHV-infected iSLK.219 cells. **(B)** Dotpvolcano plot for genes encoding unpregulated and downregulated mRNAs in iSLK.219 cells silenced for 24 hours (siControls or sieIF5B) followed by 48H DOX treatment. **(C)** Dotplots of GSEA performed using Hallmark for genes encoding unpregulated and downregulated mRNAs in iSLK.219 cells silenced for 24 hours (siControls or sieIF5B) followed by 48H DOX treatment. **(D)** Schematic of analysis performed comparing overlapping pathways from GSEA using Hallmark derived from our study described in (A) and eIF5B-depleted cells in hypoxia from Ho, et. al, 2018 *Cell Reports.* **(E)** Dotplot represents all signicantly enriched pathways for each data set described in (C) that overlap in each respective condition (KSHV infection timecourse and Hypoxic Cells).

To expand our analysis of eIF5B-mediated global transcriptional modulation, we compared host genes differentially expressed in KSHV-infected cells to those regulated by eIF5B during hypoxic stress in uninfected cells^39^ (**Figure 5D**). This analysis revealed statistically significant overlap between pathways affected by eIF5B depletion in KSHV-infected cells and those eIF5B silenced cells in hypoxic-exposed uninfected cells (**Figure 5E**). In particular, epithelial-to-mesenchymal transition (EMT) was identified as one of the most prominently activated pathways (**Figure 5E**). This notable finding points to an unanticipated role of eIF5B during infection and hypoxia, aligning with prior reports of KSHV-induced EMT and the contribution of EMT to tumor progression^5,54–56^. More striking is the enrichment of pathways associated with hypoxia and glycolysis, which further highlight the similarity of cellular remodeling between KSHV-infected and hypoxic cells (**Figure 5E**).

### Absence of eIF5B impacts infectivity and viral oncogenic capacity in MSCs

Previous studies have demonstrated that human mesenchymal stem cells (hMSCs) are both susceptible and permissive to KSHV infection, allowing them to serve as a natural reservoir for viral lytic replication^6,57–59^. To gain physiologically relevant insight into the role of eIF5B in KSHV biology, hMSCs were utilized as an additional cellular model. Of particular importance, KSHV-infected hMSCs display Kaposi’s sarcoma-like phenotypes^6,59^, which offers a robust platform to study *in vitro* cellular transformation and viral oncogenic mechanisms. We first infected hMSCs over 24 hours to evaluate eIF5B protein levels during KSHV *de novo* infection **(Figure S7)**. Expression of eIF5B increased during latency establishment, a pattern not observed by other eukaryotic initiation factors **(Figure S7)**, recapitulating findings of eIF5B accumulation in the DOX-inducible iSLK.219 system **(Figure 1)**.

To uncover eIF5B’s role in hMSCs, we silenced eIF5B and conducted *de novo* infection with rKSHV219 cultured in MSC media, which favors viral production^59^ (**Figure 6A**). Remarkably, eIF5B knockdown resulted in the loss of KSHV *de novo* infection levels in hMSCs by 50% (**Figure 6B-C & S7E**) as well as inhibiting the production of virions (**Figure 6D & S8**). Intriguingly, eIF5B silencing led to ablation of KSHV L protein, K8.1, but not IE protein ORF45 (**Figure 6E**), mirroring the observations demonstrated in iSLK.219 cells (**Figure 3**). These data further underscore the critical role of eIF5B in the synthesis of late lytic viral proteins and replication in this natural infection model.

**Figure 6:**
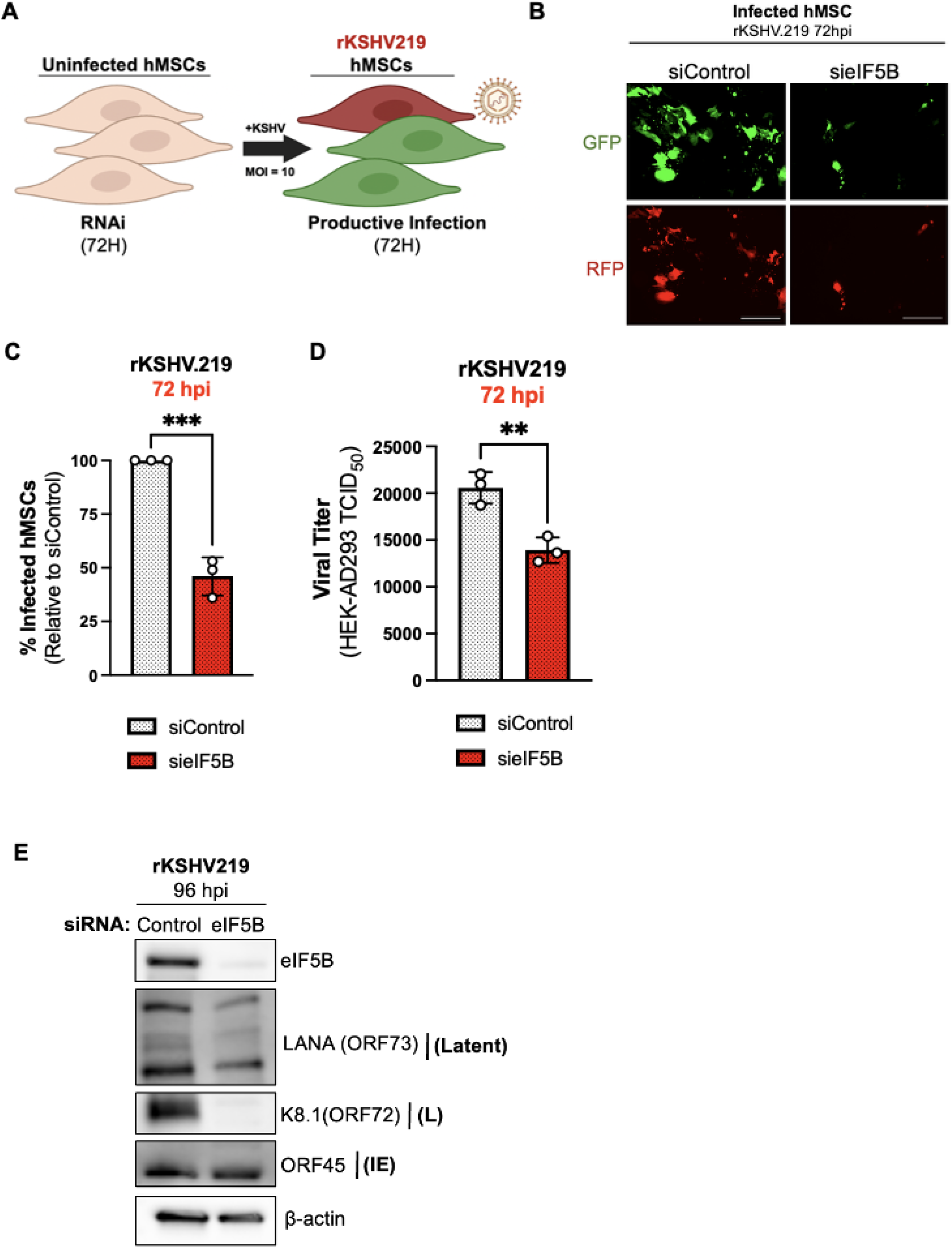
Absence of eIF5B impacts infectivity and oncogenic capacity in MSCs. **(A)** Immunoblot of eIF5B expression during KSHV-infection (MOI=10) of Human mesenchymal stem cells (hMSCs) over 24 hours. **(B)** Schematic diagram of the silencing approach for *de novo* infection of hMSCs cultured in MSC media. hMSC cells were silenced for 72H prior to infection at MOI=10 with rKSHV219. **(C)** Images of infected hMSCs that were silenced and infected as described in (A). GFP= latent KSHV infection and RFP = Lytic infection. Scale bar 100mm. **(D)** Percent of infected hMSCs after 72H of silencing sieIF5B relative to siControl measured by flow cytometry by evaluating GFP-postive cells. **(E)** Infectious virion production of siControl and sieIF5B infected-hMSCs HEK-AD293 TCID_50_ (n=3; mean ± SD; **p < 0.0060, Unpaired t-test). **(F**) Representative immunoblot of KSHV latent and lytic protein levels in siControl and sieIF5B silenced hMSCs after 96 hours of infection.

The hMSCs cultured in MSC media support KSHV infection but hinder KSHV-infected cell proliferation^6,59^. Instead, when they are grown in media abundant in angiogenic and endothelial growth factors (KS-like media), this enables persistent proliferation of infected cells, mediated by PDGFRA activation^59^. These KS-like conditions were used to assess eIF5B’s contribution to KSHV-driven oncogenic mechanisms. To accomplish this, eIF5B was silenced prior to rKSHV219 *de novo* infection, followed by culturing of infected hMSCs in KS-like media. We first assessed cellular proliferation of infected cells by using an IncuCyte Live-Cell Imaging and Analysis system which followed the growth of GFP-positive (latently infected) and RFP-positive (lytically infected) cells. Consistent with observations in the iSLK.219 model, eIF5B silencing significantly impaired establishment of rKSHV219 infection (**Figure 7A**) and suppressed the expansion of lytically reactivated cells (**Figure 7B**). Importantly, this considerable difference in proliferation was not due to eIF5B siRNA-associated effects on cell viability (**Figure S8**). Since KSHV is able to activate PDGFRA proliferative signaling through upregulation of cellular factors including the angiogenic VEGF^60,61^, we harvested the supernatants to compare the secreted VEGF levels in cells cultured in KS-like media. Reduced levels of VEGF secretion were observed in supernatants collected from infected cells lacking eIF5B (**Figure 7E**).

**Figure 7:**
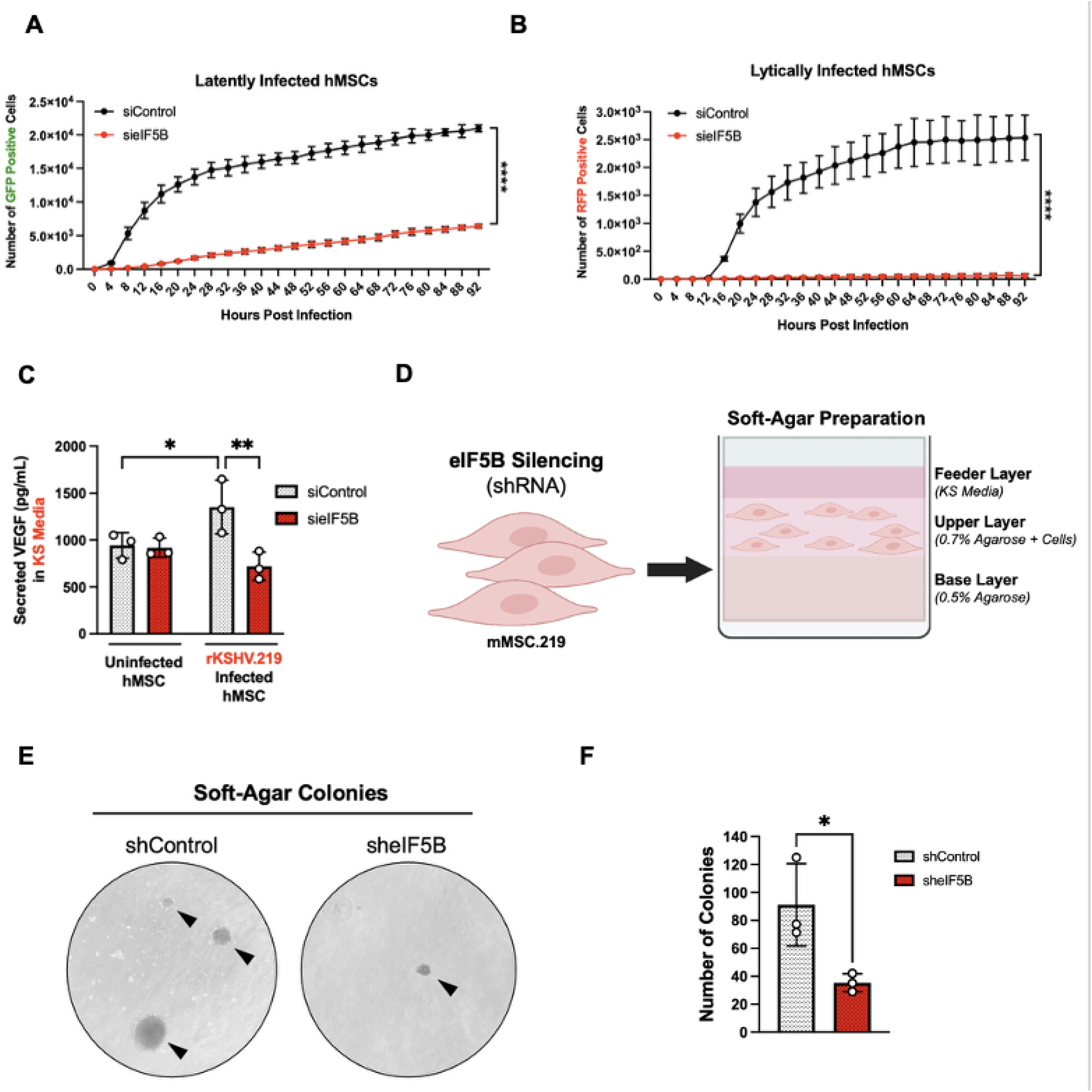
eIF5B depletion reduces viral oncogenic capacity in MSCs. **(A)** Number of latently infected hMSCs (GFP+ cells) determined by incubating silenced and infected cells as described in (A) in an IncuCyte Zoom acquiring green fluorescent images. The number of infected cells was plotted over 92 hours post-infection (n=3; mean ± SD; ****p < 0.0001, Two-way ANOVA). **(B)** Number of lytically infected hMSCs (RFP+ cells) determined as in (F) and infected cells as described in (A) in an IncuCyte Zoom acquiring red fluorescent images. The number of infected cells was plotted over 92 hours post-infection (n=3; mean ± SD; ****p < 0.0001, Two-way ANOVA). **(C)** Secreted levels of VEGF in cell-free supernatants from hMSC cells silenced and infected as described in (A). After initial spinoculation, media was changed and cells were grown in KS-like media for the 72 hours of infection. **(D)** Schematic for soft-agar colony formation assay preparation. **(E)** Representative soft-agar colonies to establish anchorage-independent cell growth of eIF5B silenced in mMSC.219 cells were grown in KS-like media and silenced using shRNA prior to being seeded in soft agar plates. **(F)** Quantification of colonies from (I) calculated using Image J (n=3; mean ± SD; **p < 0.0231, ***p < 0.0325, Unpaired T-test)

These results suggest eIF5B plays an important role in KSHV-induced cellular transformation. To test this, we sought to explore how eIF5B depletion may impair KSHV’s ability to confer malignant cellular properties, such as anchorage-independent cell growth^62^. We used mouse MSCs infected with rSKHV219 which were previously generated and shown to grow colonies in soft-agar when cultured in KS-like media^59^ (**Figure S8C**). We successfully knocked out eIF5B using shRNA (**Figure S8D-E**) in mMSC.219 cells and performed soft-agar colony formation assay (**Figure 7D**) which resulted in fewer colonies in the absence of eIF5B (**Figure 7E, 7F & S8F**). Collectively, this data supports that eIF5B is critical to KSHV infection and oncogenicity.

### Robust eIF5B staining was found in AIDS-Kaposi’s Sarcoma lesions which localize to regions of KSHV infection

The eIF4F^H^ hypoxic translation machinery is essential for protein synthesis under low oxygen conditions, including within the hypoxic cores of tumors^63^. To evaluate the relevance of eIF5B *in vivo*, we assessed the expression of eIF5B in AIDS-KS tumors **(Figure S9).** Skin biopsies representing early (patch/plaque), intermediate (plaque) and advanced (nodular) stages of AIDS-KS tumors were acquired from the AIDS Cancer Specimen Repository (ACSR) and analyzed for KSHV LANA and human eIF5B **(Figure 8A)**. We observed LANA-positivity increased progressively with advancing tumor stage, most prominently in nodular KS (**Figure S9**). While overall eIF5B expression levels did not differ significantly across early and intermediate tumor stages (**Figure S9**), eIF5B staining appeared spatially enriched within infected regions of nodular lesions (**Figure 8B**). Curiously, immunohistochemistry analysis revealed a decrease in eIF5B-positive cell numbers within KSHV-infected areas (LANA-positive) compared to uninfected regions (LANA-negative) within nodular-stage tumors (**Figure 8C**). These findings underscore a potential role for eIF5B in late-stage KSHV-tumorigenesis.

**Figure 8:**
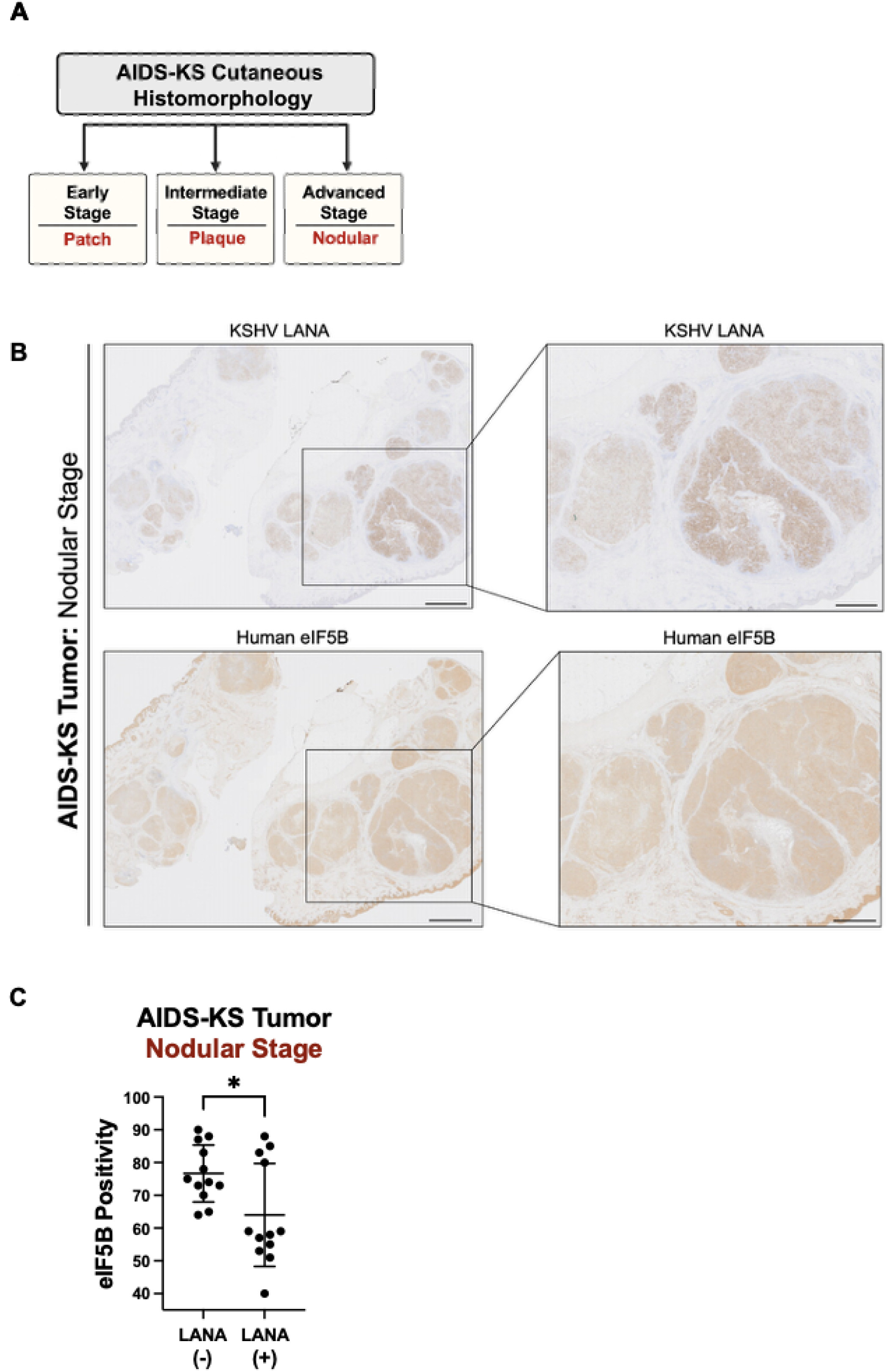
Intense eIF5B staining found in AIDS-Kaposi’s Sarcoma Lesions and localizes to regions of KSHV infection. **(A)** Schematic diagram of the staging in cutaneous AIDS-KS tumors. **(B)** Representative images of AIDS-KS nodular stage patient immunohistochemical (IHC) staining for KSHV LANA and human eIF5B. Scale bar 2mm. **(C)** Percent positive eIF5B cells in AIDS-KS nodular stage patient tumors as determined by Visiopharm imaging software. Three regions of interest (ROI) per patient that were LANA-positive were considered KSHV-infected cells and three ROI per patient that were LANA-negative were considered uninfected. (n=12, *p< 0.0275, Unpaired T-test with Welch’s Correction)

## Discussion

During viral infection, host cells use intrinsic defenses that target translation factors to suppress protein synthesis machinery^64,65^. Oncoviruses navigate these translational shutoff mechanisms by evolving strategies to limit these cellular antiviral responses to sustain persistent infections^11,14,66–68^. Here, we have identified a previously unrecognized role for eIF5B in supporting KSHV late lytic replication and driving selective viral lytic mRNA translation in normoxia when the canonical eIF2 is suppressed. These results support a translational switch from eIF2 and towards eIF5B-dependency, which recapitulates a mechanism typically engaged during hypoxic stress **(Figure 9)**.

**Figure 9:**
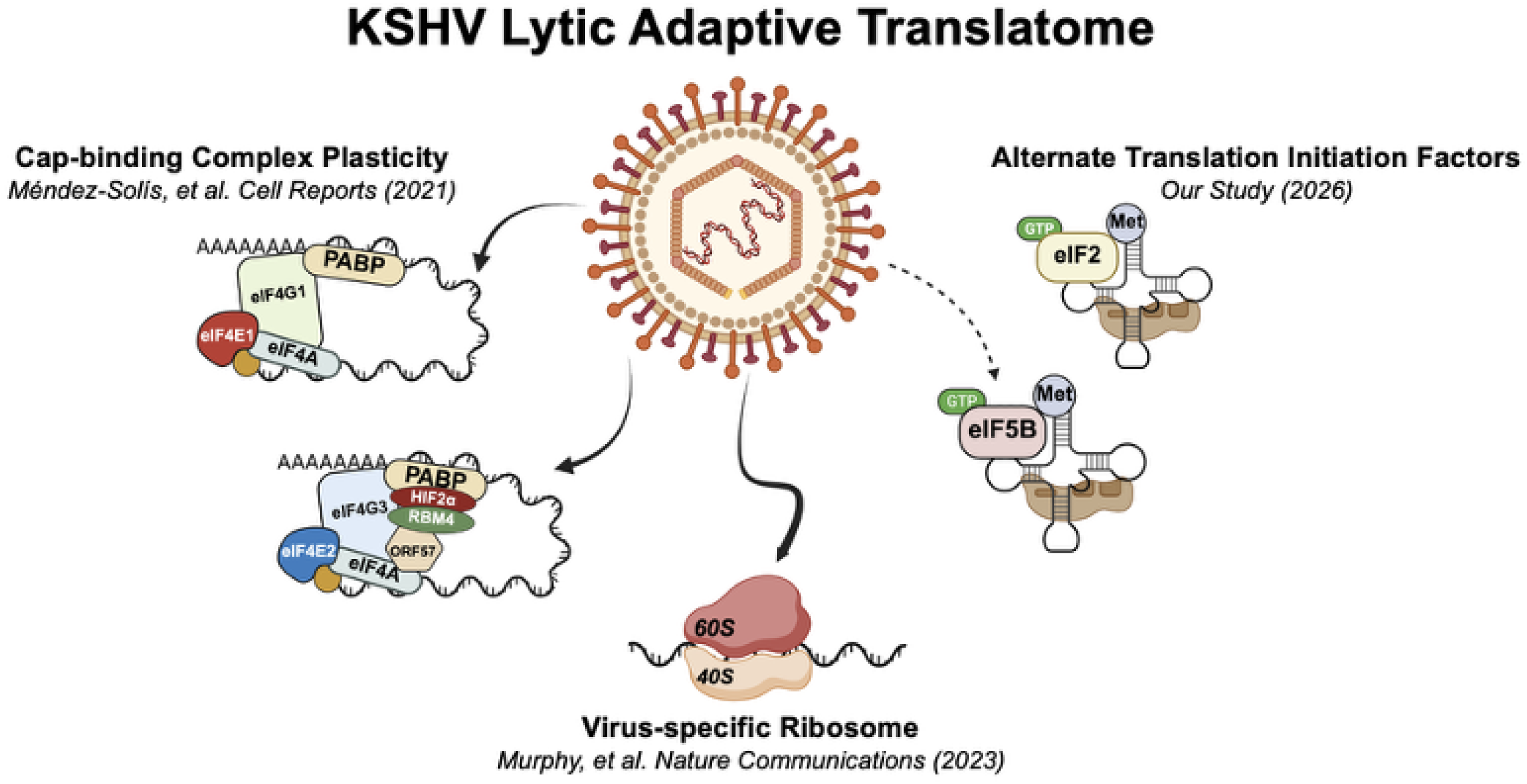
Schematic overview of updated understanding of host translation machinery utilized during KSHV infection and replication. This model demonstrates that KSHV can usurp both canonical and non-canonical host translation machinery for synthesis of lytic viral proteins and focuses on the most recent discoveries relevant to this current study. First, in Méndez-Soli’s et al. *Cell Reports* (2021), it was discovered that along with the canonical eIF4F translation initiation complex, KSHV is also able to preferentially use the hypoxic eIF4F^h^ initiation complex during normoxic conditions. This includes recruitment of the hypoxic eIF4E2 cap-binding protein, HIF2α and KSHV ORF57 to 5’ mRNA caps. Recent work has further expanded our understanding of the translation machinery used during KSHV infection. Specifically, in Murphy et al. *Nature Communications* (2023), they uncovered that KSHV mediates compositional changes to cellular ribosomes, utilizing specialized ribosomes to facilitate translation of viral mRNAs. To extend this model, our work reveals that during infection, when the canonical eIF2 is inhibited, KSHV can co-opt the hypoxic surrogate of eIF2, known as eIF5B. The utilization of these alternate initiation factors and complexes deepens our knowledge of KSHV biology and translational plasticity, demonstrating KSHV is masterful at exploiting cellular resources to ensure viral propagation.

Cells in hypoxia undergo stress-induced eIF2-inhibition through the phosphorylation of the eIF2α subunit^52,69^. Despite this, hypoxic cells engage in cap-dependent translation of mRNAs by utilizing eIF5B for the delivery of met-tRNAi^Met^ through the eIF4F^H^ complex^70^. Although under normoxic conditions, eIF5B has been shown to be dispensable to cells^70,47,71^, our data show that KSHV relies on cellular eIF5B for lytic reactivation and infectious virion production (**Figure 2**). Moreover, increased accumulation of eIF5B in polysomes during lytic infection (**Figure 3D**) reveals a dependence on this translation factor on expression of E and L viral genes (**Figures 2 & 4**). This temporal selectivity suggests that at the onset of the lytic cascade may still be reliant on residual eIF2 while suppression of eIF2 during infection leads to a progressive reliance on eIF5B, as demonstrated by accumulation of eIF5B protein levels in the late lytic stage (**Figure 1E-F**), coinciding with the surge in late viral gene expression (**Figure 3**). This data supports a model where eIF5B becomes a key component in late lytic viral translation initiation, as depletion of eIF5B did not affect ribosome density profiles in uninfected or latently infected cells (**Figures 4A & S5**). Importantly, KSHV reliance is specific to eIF5B as alternative initiation factor eIF2D silencing failed to impair polysome profiles (**Figure 4E**). Dependence on other translation factors during infection may be a shared strategy among herpesviruses as it has been reported that HCMV is also selectively dependent on an alternative translation initiation factor, eIF3d, for the synthesis of late lytic viral genes^72^.

More broadly, this study’s findings provide further evidence of stress-specific translation machinery remodeling in cellular landscapes^26,72–75^. Our RNA-sequencing analysis following knockdown of eIF5B in KSHV lytic cells uncovered a larger role for eIF5B in modulating the host transcriptome in a manner that mimics effects seen in hypoxic cells (**Figure 5**), as indicated by pathways enriched for hypoxia and glycolysis (**Figure 5**). The glycolytic switch exhibited in hypoxia has been shown to be dependent on eIF5B in uninfected cells^70,76,77^. It is tempting to propose that an oncogenic virus, like KSHV, co-opting eIF5B may unintentionally result in a step towards cellular transformation since aerobic glycolysis, commonly known as the Warburg Effect, is one of the identified hallmarks of cancer^76^. In fact, other mechanisms of KSHV-mediated glycolytic induction have been described previously^78–80^. It does not escape our attention that enrichment in immune and inflammatory pathways in our RNA-sequencing data also suggests that KSHV may exploit eIF5B to bypass host-stress and defense responses (**Figure 5**). Taken together, the overlapping transcriptional landscape between infected cells in normoxia and uninfected hypoxic cells implicates an important role in eIF5B’s contribution to the hypoxia-like environment documented during KSHV infection- a role that may facilitate immune evasion and viral oncogenesis^81,82^.

Notably, the importance of eIF5B extends beyond just the DOX-inducible iSLK.219 reactivation model. In our hMSC model for KSHV *de novo* infection, which recapitulates many aspects of KS pathobiology, eIF5B silencing impaired KSHV infectivity and virion production (**Figure 6**). Of greater significance is the suppression of viral oncogenic mechanisms, including the capacity of KSHV-infected murine MSCs to form colonies in the absence of eIF5B (**Figure 7).** We also examined eIF5B expression in AIDS-KS skin lesions from early, intermediate and advanced stages of disease progression (**Figure 8**). We observed a difference in eIF5B positivity in uninfected regions compared with adjacent KSHV-infected cells (**Figure 8**). Many cancers are associated with dysregulation of translation factors, including overexpression of certain eIFs, which points towards a potential important role for eIF5B in KSHV-driven tumorigenesis during late-stage KS^83–85^.

Therapeutically, the generation of translation inhibitors and targeting eIFs has been widely used for several malignancies^86–88^. Elucidating previously unrecognized roles for these translation initiation factors in cancer is increasingly important for revealing new vulnerabilities to therapeutically exploit. Beyond KS, eIF5B has been identified as a potent driver in translating programmed death ligand 1(PD-L1) in lung cancer, which is highly expressed in tumors and can act to suppress T-cell activity^89^. Because eIF5B is considerably overexpressed in lung adenocarcinomas and is associated with poor survival, this has implicated eIF5B as a novel intervention target in this cancer type^89^.

The evolutionary lineage of eIF5B can be traced back to the last universal common ancestor (LUCA)^70,71,90^. Our study points to an important role for this highly conserved protein during KSHV infection. By functionally replacing eIF2, eIF5B facilitates selective translation of specific KSHV transcripts. By co-opting the ancient eIF5B, KSHV can facilitate viral replication and these data add to the existing repertoire of viral strategies for translation control and specifically the hypoxic translation machinery in the context of KSHV biology^11,14,67,82^.

## EXPERIMENTAL MODELS & SUBJECT DETAILS

### KSHV Cell Lines

In this study, iSLK.KSHV219 cells, iSLK (KSHV-negative) cells and HEK-AD293 cells were cultured in Dulbecco’s modified Eagle medium (DMEM) supplemented with 10% fetal bovine serum (FBS, Gemini Bio-Products) and 1% penicillin-streptomycin (Gibco). Selection in iSLK.KSHV219 cells was conducted by culturing cells in the presence of 10μg/mL of puromycin (Gibco), 50 μg/mL of G418 (Sigma) and 50 μg/mL of Hygromycin B (Invitrogen). iSLK (KSHV-negative) cells were cultured in only 10μg/mL of puromycin (Gibco) and 50 μg/mL of G418 (Sigma).

### EBV Cell Lines

A marmoset B-Cell line infected with EBV (B95-8) was used in this study. These cells were maintained in RPMI-1640 supplemented with 10% FBS and 1% 1% penicillin-streptomycin (Gibco). To reactivate EBV form the latent to the lytic stage, 20ng/mL of TPA and 3mM of Sodium butyrate were added to each well of seeded B95-8 cells.

### Primary Human Cells

Primary human mesenchymal stem cells (hMSCs) were isolated as previously described (Gomes et al. 2013), To maintain hMSCs in culture, this study used MSC or KS-like media as described in previous studies (Gomes 2013, Naipauer 2019, Mendez 2021, Lacunza 2024). MSC media: Minimum essential medium (αMEM/Invitrogen) supplemented with 20% FBS (Atlanta Biologicals) and 1% penicillin-streptomycin. KS-like media: Dulbecco’s modified eagle medium (DMEM/Corning) containing 30% FBS (Gemini Bio-Products), 0.2mg/mL Endothelial Cell Growth Factor (ECGF) (ReliaTech), 0.2 μg/mL Endothelial Cell Growth Supplement (ECGS) (Sigma-Aldrich), 1.2 μg/mL heparin (Sigma-Aldrich), insulin/transferrin/selenium (Invitrogen), 1% penicillin-streptomycin (GIBCO) and MEM vitamin (VWR Scientific) at 37°C and 5% CO2.

### PDGFRA-positive mouse MSCs

Mouse mesenchymal stem cells (mMSCs) were isolated and infected with rKSHV.219 (mMSC.219) as previously described (Naipauer 2019). To maintain these PDGFRA-positive and KSHV-infected murine cells, they were cultured in MSC media, made up of MEM alpha media supplemented with 20% heat inactivated fetal bovine serum (FBS) and 1% penicillin-streptomycin. Following growth in MSC media, mMSC.219 cells were cultured in KS-like media as described above.

### Human Tissue Samples

For this study, all human specimens were provided by the AIDS and Cancer Specimen Resource (ACSR), funded by the U.S. National Cancer Institute. Tissue samples were from 10 human patients with Kaposi’s Sarcoma and were collected in accordance with the Federal human subjects regulations, 45 C.F.R., Part 46. Samples were selected based on pathological evaluation of 60% or more tumor content. All tissue samples from human cases have been de-identified in accordance with the Health Insurance Portability and Accountability Act (HIPAAA).

## METHOD DETAILS

### Virus Production

To prepare viral inoculum, iSLK.KSHV219 cells harboring the recombinant rKSHV219 virus were used (Myoung and Ganem 2011). KSHV virus was concentrated as previously described (Naipauer PLoS Pathogens 2019 & Rosario et al 2018). Briefly, rKSHV219 virus from iSLK cells were induced at 50-60% confluency by the addition of doxycycline (1μg/uL, Sigma-Aldrich) and sodium butyrate (1mM) for 4 days total. Supernatants from induced iSLK.KSHV219 cell cultures were collected and filtered through 0.45-μm filters followed by ultracentrifugation at 25,000 rpm for one hour and a half at 4°C. After centrifugation the virus pellet was resuspended in DMEM base medium (COMPANY) without supplementation. Pellets were pooled and aliquots were stored at −80°C. Virus titer was calculated by infecting HEK-AD293 cells with serial dilutions of rKSHV219 virus. At 72 hours post-infection, GFP+ cells were then counted by flow cytometry to determine viral titer.

### Virus concentration using PEG-it^TM^ virus precipitation solution

KSHV virus was also produced using the PEG-it^TM^ virus precipitation solution (Systems Biosciences, Cat. No. LV810A-1). Infectious rKSHV219 virus from iSLK cells (Myoung and Ganem 2011) were induced at 50-60% confluency by the addition of doxycycline (1μg/uL, Sigma-Aldrich) and sodium butyrate (1mM) for 4 days total. Supernatants from induced iSLK.KSHV219 cells were collected and filtered through 0.45-μm filters followed by adding 1 volume of PEG-it^TM^ virus precipitation solution to 4 volumes of viral supernatant and gently mixed at 4°C overnight. Viral pellets were collected and washed once prior to resuspension in DMEM base medium (COMPANY) without supplementation. Pellets were pooled and aliquots were stored at −80°C. Virus titer was calculated by infecting HEK-AD293 cells with serial dilutions of rKSHV219 virus. At 72 hours post-infection, GFP+ cells were counted by flow cytometry to determine viral titer.

### RNAi delivery through siRNA transfection

All cells were transiently transfected with small interfering RNA (siRNA) for 24 hours prior to viral reactivation or 72 hours before *de novo* infection at a final concentration of 25 pmol per well using Lipofectamine RNAiMAX reagent (Life Technologies). Each siRNA used in this study was purchased from GE Dharmacon.

### Lentiviral transduction & shRNA gene silencing

Lentiviral vectors were derived following the transfection of HEK-293T cells with packging plasmids pMD2-G and Pax-2 (COMPANY) using FuGene (Company). After 72 hours post transfection, supernatant containing lentivurs particles were harvested and cellular debris was removed following centrifugation at 2500 rpm for 20 minutes and filtration using 0.45-μm filters. To concentrate lentivirus, 1 volume of PEG-it^TM^ virus precipitation solution was added 4 volumes of viral supernatant and gently mixed at 4°C overnight (System Biosciences).

### KSHV lytic induction & HEK-AD293 TCID_50_

Following 24 hours of siRNA treatment, iSLK.KSHV219 cells were induced upon reaching 60% confluency using doxycycline (1μg/uL, Sigma-Aldrich). After 48-72 hours of induction, cell-free virus containing supernatants were harvested and stored at −80°C until virus titer determination. HEK-AD293 cells were seeded at 8 × 10^4^ into each well in 12-well plate 24 hours prior to infection. The following day, HEK-AD293 cells were pre-treated with 8μg/mL of polybrene (Millipore, Cat No. TR-1003-G) for 30 minutes at 37°C. Thawed viral supernatants were filtered through 0.45-μm filters and used to *de novo* infect these pretreated cells by spinoculation at 700 x *g* for 60 minutes at 30°C. Infected cells were incubated at 37°C for 72 hours and then infectious virion production was measured by counting GFP-positive HEK-AD293 cells using flow cytometry to determine the 50% tissue culture infective dose (TCID_50_) where 1 GFP+ HEK-AD293 = 1 Infectious KSHV virion.

### rKSHV219 *de novo* infection of hMSCs

Human MSCs were transiently silenced for 72 hours using siRNA. Then, silenced hMSCs were pre-treated with 8μg/mL of polybrene (Millipore, Cat No. TR-1003-G) for 30 minutes at 37°C. Pre-treated hMSCs were then infected with rKSHV219 at multiplicity of infection (MOI) of 10 by spinoculation at 700 x *g* for 60 minutes at 30°C. To determine infectious virus titer, virions in the supernatants were harvested 72 hours after infection and used for *de novo* infection of HEK-AD293 cells as described above.

### Western Blotting

Radioimmunoprecipitation assay lysis buffer (RIPA, ThermoScientific) containing phosphatase and protease inhibitors (Sigma) was used to obtain protein lysates. Lysates collected were sonicated and centrifuged at 10,000 rpm for 10 minutes at 4°C to remove genomic DNA. Protein concentration was measured by using bicinchoninic acid (BCA) protein assay (ThermoScientific) prior to the addition of Laemmli buffer (BioRad) containing β-mercaptoethanol (Millipore) to each sample. Equal amounts of protein was loaded and resolved in SDS-PAGE gels (BioRad). Each gel was then transferred to a polyvinylidene difluoride (PVDF) membrane (BioRad) and blocked with 5% BSA (Sigma) for 1 hour for reduction of non-specific binding. Then, membranes were incubated with primary antibodies diluted in 5% BSA overnight at 4°C. Protein bands were visualized using SuperSignal West Pico, Femto or Alto PLUS Chemiluminescent Substrate (ThermoScientific). For extracted protein from polysome profiles, equal volumes of lysate were added to each lane and processed as described above. The intensity of each western blot band was measured using ImageJ software. Uncropped data for all representative immunoblots are provided (**Supplementary Figure 9**).

### Episome Copy Number

To measure episome copy number we silenced latent and reactivated iSLK.KSHV219 for 24-72 hours. Cells were then harvested according to the PureLink® Genomic purification kit (ThermoFisher) protocol. Following DNA extraction, KSHV copy number in 30ng of DNA was determined using the Applied Biosystems QuantStudio Absolute Q Digital PCR(dPCR) System (ThermoFisher).

### Cell Viability

To determine cell viability, cells were washed with PBS and harvested using 0.05% trypsin (Gibco) for mesenchymal stem cells or 0.25% trypsin (Gibco) for all other cells for 5 minutes in 37°C. They were then resuspended in media and mixed 1:1 with 0.4% trypan blue solution (Sigma). Viability was calculated as the percentage of unstained or viable cells and number of live cells using an automatic cell counter (BioRad).

### Real-time quantitative PCR (RT-qPCR)

RNA was extracted from cells using RLT buffer (RNeasy Kit Qiagen) containing β-mercaptoethanol (Millipore) or by phenol-chloroform extraction (Fisher Scientific) with ethanol precipitation. To remove DNA, each sample was treated with RNase-Free DNase (Qiagen) on columns for 25 minutes or with amplification-grade DNAase I (Sigma) for 15 minutes at room temperature. RNA was reverse transcribed into cDNA using lmProm-II Reverse Transcriptase (Promega) as directed by the manufacturer’s protocol. Viral and cell host mRNA’s were amplified using specific primers (Table S1) diluted in SYBR Green PCR master mix (Quanta Biosciences). For detection, this study used the PowerGene 9600 Plus Real-time PCR system (ATILA BioSystems). Non-reverse transcriptase and water controls were used to confirm the absence of viral DNA and contamination in each sample. β-Actin was used as a housekeeping gene to perform ΔΔCT method. The expression of host and viral genes was normalized to β-Actin CT values and the difference between the host and viral CT values with actin CT was considered the ΔCT value. The obtained ΔCT values were normalized to a given control sample (ΔΔCT) and the fold change was calculated using the 2^-ΔΔCT^ formula. For analysis of viral genes obtained after polysome profiling, the CT values of each fraction were normalized to the input RNA of each viral gene (ΔCT). For oligosome and 2fraction (ΔΔCT value). The fold change was obtained using the 2^-ΔΔCT^ formula.

### Flow Cytometry

In order to measure the amount of infected (GFP+) cells for the HEK-AD293 cell line, latently infected (GFP+) or lytically reactivated (GFP+/RFP+) in iSLK.KSHV219 or hMSCs, cells were washed two times with 1X PBS and fixed in 4% paraformaldehyde. For each sample, 10,000 cells were recorded. Flow cytometry analysis was performed using a Becton-Dickinson LSR analyzer (BDBiosciences).

### Puromycin Incorporation Assay

B95-8 EBV cell lines were treated with 101μg/mL puromycin for 10 minutes prior to harvest. To determine puromycin incorporation into nascent proteins, this study used western blotting with an antipuromycin antibody. For quantitative analysis, the total signal of antipuromycin in the Western blot was normalized to the total protein signal captured using SWIFT membrane stain (Biosciences).

### Ribosome Density Fractionation

Polyribosome fractionations were performed in iSLK.KSHV219 cells as previously described (Mendez). Briefly, iSLK.KSHV 219 cells were silenced as outline above and grown in 15-cm dishes to 95% confluency grown. Cells were treated with 0.2mg/mL cycloheximide (VWR) for 10 minutes prior to harvesting cells in order to immobilize actively translating ribosomes. RNA lysis bugger [15 mM Tris-HCL (pH= 7.4), 15mM MgCl_2_, 0.3M NaCl, 1% Triton X-100, 0.2mg/mL cycloheximide and 200 units/ml RNaseOut (Invitrogen)] was used to collected immobilized translating ribosomes. Equal number of RNA from each group was loaded onto a 10-50% sucrose density gradient. The gradients were sedimented by centrifugation at 39,000 rpm for 90 minutes at 4°C and fractionated using Brandel BR-188 density gradient fractionation system. Peakchart software (Brandel) was used to visualize polysome profiles. Each fraction was treated with proteinase K (Ambion) to obtain mRNAs using phenol-chloroform extraction with ethanol precipitation.

### Cell Proliferation Assay (IncuCyte)

Proliferation was measured as previously described (Naipauer 2019 and Mendez 2021) Briefly, human MSCs were plated in 12-well plates at 25,000 cells/well in 3 replicates and silenced for 72 hours in MSC media. Following this silencing, hMSCs were infected with rKSHV219 at an MOI of 10 and then infected cells were incubated in an Incucyte Zoom (Essen Bioscience) to acquire green and red fluorescence images at 10X magnification every 4 hours. Incucyte Zoom software was used to process and analyze the results.

### Enzyme-linked Immunosorbent Assay (ELISA)

After 72 hours of infection with rkSHV219 or mock infection, supernatants from uninfected and infected hMSCs cultured KS media were harvested. A human VEGF-A ELISA kit was used determine the concentration of VEGF-A in cell-free supernatants according to the manufacturer’s instructions (Cusabio). The absorbance was measured by using a plate reader at 450nm (Molecular Devices) and VEGF-A quantity was calculated using a standard curve.

### Soft-Agar Colony Formation Assay

Colony formation assay was conducted as previously described (Naipauer 2019). Briefly, base agar was made by combining melted 1% agar with KS-like medium to give a 0.5% agar/1X KS-like medium solution followed by addition of the 1.5mL of this solution to each well of a 12-well plate. Melted 0.6% agarose was mixed with 2X KS-like media and mMSC.219 cells to seed 2,500 cells per well in a 12-well plate. Cells were fed KS-like media every 2-3 days. Colonies were counted and photographed after 4 weeks and GFP+ colonies were confirmed by microscopy to ensure KSHV infection.

### Immunohistochemistry (IHC) staining of KS tumors

The 5-micron FFPE tissues were pre- baked at 60°C for 30 minutes to improve slide adhesion. The Immunohistochemistry (IHC) staining was performed on the Leica Bond RX automated platform. Sections were dewaxed and rehydrated before performing the Epitope Retrieval for 20 minutes at 100°C (Leica Bond ER1 Ref#AR9961) to expose the target. Then the tissues were blocked with 10% Normal Goat in PBS and incubated with the primary antibody at the optimal dilution for 30 minutes at room temperature (EIF5B, Polyclonal, Proteintech 13527-1-AP; LANA (KSHV) (13B10), Leica Biosystems PA0050-U). It was used Bond Polymer Refine as detection system (Leica, Ref#DS9800) with the chromogen DAB precipitate at the antibody binding site followed by counterstaining with light Hematoxylin. The stained slides were dehydrated and mounted with micromount using the automated stainer-cover slipper (Leica ST5020, CV5030).

### Bioinformatics and statistical analyses of RNA-seq data

RNA-seq paired-end reads were mapped onto the human genome GRCh38.p10 using aligner Start27.11b^91^ by performing the pipeline nextflow nf-core/rnaseq^92^. Adaptors and quality were trimmed and the biased methylation positions for directional and non-directional were removed by performing Trim Galore^93^. Tool BBSplit was used to remove genome contaminants and bin reads by mapping to multiple reference genomes simultaneously. The rRNA sequences were removed by using SortMeRNA^94^. We used samtools to sort bam files and Salmon to quantify the expression of transcripts, picard MarkDuplicates to mark duplicate reads. After completing nextflow pipeline, a gene count matrix was generated and output. In the R environment, the data were normalized by utilizing variance stabilizing transformation of DSEq2^95^ from the raw gene count data and used for doing principal component analysis (PCA) for classification of samples and making PCA plot. We also performed DESEq2 to normalize the count data. In the normalized data, we performed DESEq2 to identify genes differentially expressed (DE) in comparisons (siControl vs sieIF5B in 0h, 12h, 24h, 48h, and 72h; 0h vs 12h, 0h vs 24h, 0h vs 48h and 0h vs 72 in sieIF5B) at cutoff p-value adjusted by Benjamin-Hochberg procedure < 0.05. We used heatmap to show differential expressions of DE genes and utilized DE genes to analyze pathways in hallmark (category H), and canonical pathways including KGG, BioCarta, reactome and WikiPathways (category C2) from R fgsea^96^. GSEAplot was made by using GSEA with hallmark gene set. The pathways were selected by setting adj-pvalue (FDR) < 0.05, barplots were made by using NES of pathways, and dotplots were made by using NES and FDR in R environment. We downloaded bam data (access number: SRP110475) from SRA: SRP110475 - SRA - NCBI, converted bam files to fastq files and re-performed nextflow-nf-core/rnaseq to obtain count data (TPM data) of genes. We chose to compare 0h, 12h, 24h,48h and 72h in sieIF5B with 9-Hypo-eIF5B-Poly. We chose up-regulated genes by setting log2(Hypo_9_eIF5B_Poly/ Hypo_6_Poly) > 1 and down-regulated genes using log2(Hypo_9_eIF5B_Poly/ Hypo_6_Poly) <-1. These genes selected were used to perform pathway analysis in in hallmark and canonical pathways. These pathways were compared to those obtained from our RNA seq data.

### Statistical Analysis and Quantification

Experiments in this study were conducted at least three independent times unless otherwise stated. All values were imported into GraphPad Prism 10 software for appropriate statistical analysis, e.g., Student’s t test or 2-way ANOVA. For all experiments in this study, statistical significance was defined by *p* < 0.05. Densitometry analyses of immunoblot bands were performed using ImageJ online software as described in the figure legends.

### In Memoriam

Dr. Enrique A. Mesri, inspiring scientist, dedicated mentor and dear friend, who sadly passed away during the course of this study. This work is dedicated in honor of his impact in the fields of KSHV and viral oncology and his deep commitment to training a new generation of researchers.

## Acknowledgments

We would like to thank the Flow Cytometry Core Facility for all assistance with flow cytometry. Use of the IncuCyte live-cell analysis was provided by support of the shared equipment funds provided by the Sylvester Comprehensive Cancer Center. We would like to express gratitude to Dr. Jonathan Schatz and Tyler Cunningham for guidance and use of their laboratory’s ribosome density fractionator and Dr. Juan Carlos Ramos for sharing B95-8 EBV cell line. To CD Genomics, for their assistance in ribosome density fractionation. This work was supported in part by the National Institutes of Health’s National Cancer Institute (NCI), grant CA136387 to E.A.M. and N.S. and NCI grant CA250072 to E.A.M., S.R., and N.S., National Cancer Institute (NCI) diversity supplement grant 3R01CA136387-14S1 to C.M., National Institute of Allergy and Infectious Diseases (NIAID) of the National Institutes of Health grant AI165337 to N.S. and the Florida Department of Health through the Bankhead-Coley Cancer Research Program grant 21B10 to N.S.., The D.C. Center for AIDS Research (CFAR) also provided subaward grant 5P30AI117970-07 to C.M. Specimens were provided by the AIDS and Cancer Specimen Resource (ACSR), funded by the U.S. National Cancer Institute.

## Author Contributions

Conceptualization, C.M., E.A.M. S.L. and N.S.; Methodology C.M., O.M.S, R.K., J.N., A.A., E.A.M., S.L. and N.S.; Investigation C.M., O.M.S, R.K., J.N., A.A., N.R.O., C.D.N., T.M.D.T.; Resources C.M., E.A.M. S.L., J.H.M., S.R. and N.S.; Data Analysis C.M., O.M.S, R.K., J.N., A.A., E.A.M., S.L. and N.S.; Writing-Original Draft, C.M. and N.S.; Writing-Review & Editing, C.M., S.L. and N.S.; Bioinformatic Analysis, Y.T.; Histopathological Specimen Immunohistochemistry (IHC) of AIDS-KS Lesions, Y.M.; IHC Examination and Analysis, E.K.; Supervision, E.A.M. and N.S.

## Data Availability Statement

Further information and requests for resources or reagents can be provided upon reasonable request. The bioinformatic data that support these findings are openly available at the following temporary site which is the submitted accession number to NCBI: SUB16007332.

---

**Table S1:**
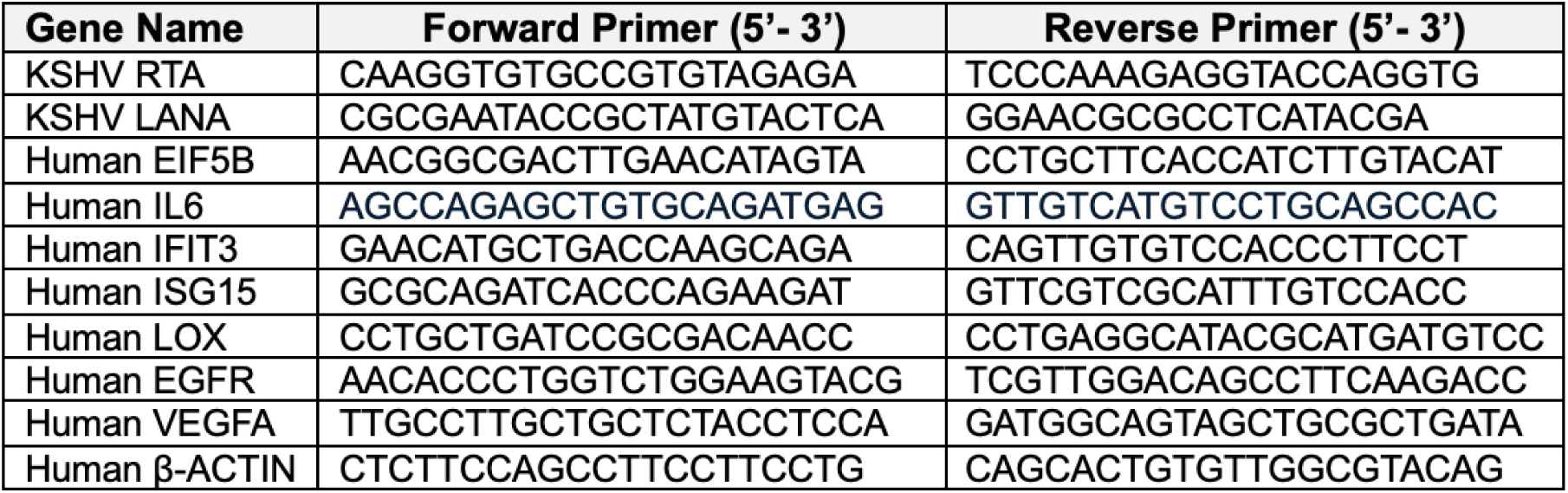
Oligonucleotide Sequences.

**Table S2:**
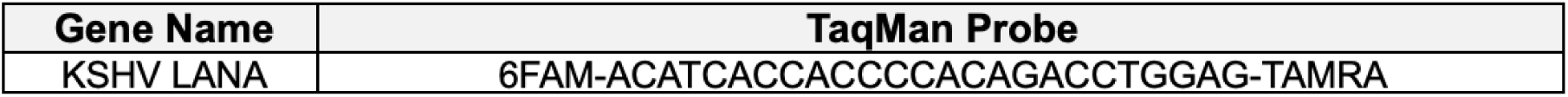
Probe Design.

**Figure S1: Eukaryotic initiation factors protein levels do not increase during KSHV infection, related to Figure 1**

**(A)** Immunoblot of eIF5B, HIF1α and HIF2α protein levels in uninfected iSLK cells after 24 hours in hypoxia (1% O_2_). **(B)** Schematic diagram of the canonical eIF4F translation inititation complex and the alterantive eIF4F^H^ complex used by cells under hypoxic stress. **(C-D)** Immunoblot of selected eIF4F translation factor protein levels at (0H) and after (24-72H) DOX treatment in iSLK.219 cells cultured in normoxia. Densitometric analysis determined by using Image J.

**Figure S2: Flow cytometry gating strategy (related to Figure 1) and knockdown of eIF5B does not affect the cell viability of uninfected iSLKs and iSLK.219 cells, related to Figure 2**

**(A)** Representative flow cytometry plots to determine percentage of reactivation 48H post-dox of sieIF5B relative to siControl. RFP expression was considered the reactivation marker and was measured using flow cytometry. Uninfected iSLK cells were used as a negative control. iSLK.BAC16 cells were used as a GFP+ control and iSLK.RGB cells were used as an RFP+ control. **(B)** Representative flow cytometry plots to determine infectious virus titer from siControl and sieIF5B iSLK.219 cell free supernatants i. cells post-reactivation Infected HEK-AD293 were determined by GFP+. Uninfected iSLK cells were used as a negative control. iSLK.BAC16 cells were used as a GFP+ control. **(C)** Percent live cells after 72 hours silencing (siControl or sieIF5B) in unreactivated iSLK.219 cells (n=3, ns= no significance, Unpaired T-test). **(D)** Number of live cells after 24- 72 hours silencing (siControl or sieIF5B) in unreactivated iSLK.219 cells (n=3, ns= no significance, Unpaired T-test). **(E)** Percent live cells after 72 hours silencing (siControl or sieIF5B) in uninfected iSLK cells (n=3, ns= no significance, Unpaired T-test). **(F)** RTA mRNA levels in DOX-inducible uninfected iSLK cells relative to actin was measured by qRT-PCR n=3, ns= no significance, Unpaired T-test with Welch’s Correction).

**Figure S3: eIF5B silencing does not impair latent KSHV infection, related to Figure 3**

**(A)** Images of iSLK.219 cells after silencing for 24-72 hours. Scale bar 100μm. **(B)** Number of latently infected cells (GFP+) from (A) after 72 hours of silencing (siControl or sieIF5B) in unreactivated iSLK.219 cells as determined by flow cytometry. (n=3, ns= no significance, Two-way ANOVA with Sidak’s multiple comparisons test). **(C)** Episome copy number of KSHV genomes in unreactivated iSLK.219 cells in either siControl or sieIF5B treatment from 24-72hrs. Data presented is the number of copies in 30ng of DNA. Copy numbers were determined using digital PCR (dPCR) (n=3, ns= no significance, Two-way ANOVA with Sidak’s multiple comparisons test). **(D)** Images of iSLK.219 cells that were silenced for 24 hours (siControl or sieIF5B). Images indicate before (0H) and after (48-72H) DOX treatment in normoxia. Scale bar 100μm. **(E)** Number of latently infected cells (GFP+) from (D) after 72 hours of silencing (siControl or sieIF5B) in DOX-induced iSLK.219 cells as determined by flow cytometry. (n=3, ns= no significance, Two-way ANOVA with Sidak’s multiple comparisons test). **(F)** Representative flow cytometry plots to determine number of latently infected cells (GFP+) from (D) treated with siControl or sieIF5B iSLK.219. Uninfected iSLK cells were used as a negative control. iSLK.BAC16 cells were used as a GFP+ control. **(G)** Representative immunoblot of KSHV LANA (latent) protein levels in siControl and sieIF5B silenced cells 48H after DOX induction. **(H)** Quantification of (G) calculated using Image J (n=3, ns= no significance, Two-way ANOVA with Sidak’s multiple comparisons test).

**Figure S4: Depletion of eIF5B does not impair the polysome profile in latently infected cells, Related to Figure 4**

**(A)** Immunoblot validating siRNA knockdown of eIF5B after silencing reactivated uninfected iSLK cells for 48 hours. **(B)** Immunoblot validating siRNA knockdown of eIF2D after silencing reactivated iSLK.219 cells for 48 hours. **(C)** Ribosome density profiles of unreactivated iSLK.219 cells silenced (siControl or sieIF5B) for 48 hours. Silenced cells were treated with cyclohexamide and lysates were sedimented through sucrose gradients and fractionated. Ribosomal subunits (40S and 60S), monosomes (80S), oligosomes (oligo) and polysomes (poly) are indicated.The table shows the area under the curve (AUC) for each fraction.

**Figure S5: Silencing of eIF5B in EBV-infected cells does not inhibit global protein synthesis, Related to Figure 4**

**(A)** Total protein of EBV-infected B95-8 cells (unreactivated or reactivated with 4uL/mL PMA and 3mM NaBut) after 72 hours silencing (siControl or sieIF5B). Membranes were washed in Total Swift tain and images taken prior to blocking membranes for immunoblot anaysis. **(B)** EBV-infected B95-8 cells (unreactivated or reactivated with 4uL/mL PMA and 3mM NaBut) after 72 hours silencing (siControl or sieIF5B). 10ug/mL of puromycin was added to all samples for 10 minutes prior to harvest and quantified relative to the total protein from (A).

**Figure S6: Validation of eIF5B knockdown in iSLK.219 cells for RNA-sequencing analysis, Related to Figure 5**

**(A)** Fold change of eIF5B from siControl and sieIF5B knockdown in iSLK.219 cells after 48H of DOX treatment (n=3; mean ± SD; *p < 0.00149, Unpaired T-test with Welch’s Correction). **(B)** Fold change of cell host mRNAs validating differentially expressed genes from Figure 5B-C (n=3; mean ± SD; ****p < 0.0001, One-Way ANOVA with Dunnett’s Multiple Comparison Test). **(C)** Fold change of cell host mRNAs validating of differentially expressed genes from Figure 5B-C (n=3; mean ± SD; ****p < 0.0001, One-Way ANOVA with Dunnett’s Multiple Comparison Test)

**Figure S7: eIF5B does not hinder the proliferation of uninfected hMSCs, Related to Figure 6**

**(A)** Images of uninfected hMSCS cells in MSC media to confirm rKSHV219 infection in infected MSCs. Scale bar 100μm. **(B)** Immunoblot of LANA expression during KSHV-infection (MOI=10) of Human mesenchymal stem cells (hMSCs) over 24 hours, confirming latency establishment in hMSCs. **(C)** Immunoblot of eIF5B expression during KSHV-infection (MOI=10) of Human mesenchymal stem cells (hMSCs) over 24 hours. **(D)** Immunoblot of eIF4E1 expression during KSHV-infection (MOI=10) of Human mesenchymal stem cells (hMSCs) over 24 hours. **(E)** Representative flow cytometry plots to determine number of infected hMSCs (GFP+). hMSCs were silenced for 72H and inoculated with rKSHV219 for 72H in MSC media. Uninfected hMSC cells were used as a negative control. iSLK.BAC16 infected hMSC cells were used as a GFP+ control

**Figure S8: eIF5B silencing in hMSCs (Related to Figure 7) and confirmation of KSHV infection in mouse MSCs (mMSC) in both cell lines and colonies formed in soft-agar plates (Related to Figure 6)**

**(A)** Representative flow cytometry plots to determine infectious virus titer from siControl and sieIF5B hMSC cell free supernatants. hMSCs were silenced for 72H and inoculated with rKSHV219 for 72H in MSC media. Infected HEK-AD293 were determined by GFP+ indication. Uninfected hMSC cells were used as a negative control. iSLK.BAC16 infected hMSC cells were used as a GFP+ control. **(B)** Number of live cells after 96 hours silencing (siControl or sieIF5B) in uninfected hMSCs cultured in either MSC or KS-like media (n=3, ns= no significance, Unpaired T-test). **(C)** Images of uninfected mMSCS cells and rKSHV129 mMSCs in KS-like media to confirm rKSHV219 infection. Scale bar 100μm. **(D)** Immunoblot validating shRNA knockdown of eIF5B after silencing mMSC.219 cells for 72 hours. **(E)** Number of live cells after 72 hours shRNAsilencing (siControl or sieIF5B) in mMSC.219 cells cultured in KS-like media (n=3, ns= no significance, Unpaired T-test). **(F)** Images of mMSC.219 colonies in soft-agar following shRNA silencing and grown in KS-like media. Images were taken after 4 weeks to confirm that these colonies were KSHV-positive. Scale bar 100μm.

**Figure S9: KSHV LANA and human eIF5B intensity in AIDS-KS skin lesions, related to Figure 7**

**(A)** AIDS-KS tumor sample immunohistochemically stained for KSHV LANA demonstrating Visiopharm software analysis and quantification parameters. Yellow indicates light staining (Histology Score +1), Orange indicates moderate staining (Histology Score +2), Red indicates strong/dark staining (Histology Score +3), Blue indicates negative stained nuclei. **(B)** Percent positive LANA cells of whole tumors from all 3 stages of AIDS-KS cutaneous lesions. Samples derived from nine HIV-positive patients provided by the AIDS Cancer Specimen Repository (ACSR) (n= 2 or 3 patients; mean ± SD; *p < 0.0289, One-way ANOVA). **(C)** Representative images of AIDS-KS patch/plaque stage patient immunohistochemical (IHC) staining for KSHV LANA and human eIF5B. Scale bar 2mm. **(D)** Percent positive eIF5B cells in AIDS-KS patch/plaque as determined by Visiopharm imaging software. Three regions of interest (ROI) per patient that were LANA-positive were considered KSHV-infected cells and three regions that were LANA-negative were considered uninfected. (n=6, ns= no significance, Unpaired T-test). **(E)** Representative images of AIDS-KS plaque stage patient immunohistochemical (IHC) staining for KSHV LANA and human eIF5B. Scale bar 2mm. **(F)** Percent positive eIF5B cells in AIDS-KS plaque as determined by Visiopharm imaging software. Three regions of interest (ROI) per patient that were LANA-positive were considered KSHV-infected cells and three regions that were LANA-negative were considered uninfected. (n=12, ns= no significance, Unpaired T-test)

**Figure S10: Uncropped immunoblots.** Protein ladder positions indicated in blue and cropped regions indicated by orange boxes.

